# Conformational Selection in Ligand Recognition by the First Tudor Domain of PHF20L1

**DOI:** 10.1101/2020.04.29.069500

**Authors:** Mengqi Lv, Jia Gao, Mingwei Li, Rongsheng Ma, Fudong Li, Yaqian Liu, Mingqing Liu, Jiahai Zhang, Xuebiao Yao, Jihui Wu, Yunyu Shi, Yajun Tang, Yueyin Pan, Zhiyong Zhang, Ke Ruan

## Abstract

The first Tudor domain of PHF20L1 (PHF20L1 Tudor1) recognizes both histone methylation and non-histone methylation to play versatile roles, e.g., PHF20L1 Tudor1 binds to the oncogenic target DNA (cytosine-5) methyltransferase 1 (DNMT1) to prevent it from degradation. However, the crystal structure of the PHF20 Tudor domain, a homolog of PHF20L1, reveals a closed aromatic cage of the Tudor domain. It is thus highly desirable to interrogate the ligand-recognition mechanism of PHF20L1 Tudor1, which will in turn validate the potential druggability of this target. Here, we solved the crystal structure of the free form PHF20L1 Tudor1, which adopts the closed conformation similar to PHF20. NMR relaxation dispersion and molecular dynamics (MD) simulations suggest a pre-existing low-population conformation with a remarkable rearrangement of aromatic cage residues. Such structural rearrangement is further revealed by the crystal structures of PHF20L1 Tudor1 in complex with the lysine 142 methylated (K142^me1^) DNMT1, and a small molecule cosolvent 2-(N-morpholino)ethanesulfonic acid (MES), respectively. This result thus ignites interest in the discovery of small molecule inhibitors against PHF20L1 Tudor1. The hit identified from NMR fragment-based screening protrudes into the same open form aromatic cage of PHF20L1 Tudor1, and blocks the interaction between PHF20L1 Tudor1 and methylated DNMT1. Further free form crystal structures of key mutants reveal one open form and one closed form aromatic cage, which is energetically trapped observed in the NMR relaxation dispersion and MD simulations. The binding of DNMT1 with PHF20L1 Tudor1 mutants was also recapitulated in cancer cells. The mutagenesis thus alters the structure, dynamics and eventually the function of PHF20L1 Tudor1. Our results demonstrate that PHF20L1 Tudor1 utilizes the same conformational selection mechanism to recognize ligands, regardless of whether it is a natural substrate or a small molecule identified from fragment-based screening. Albeit at a low population, the pre-existing ligand-binding conformation shall shift the paradigm in the druggability assessment of a dynamic protein, even though it may lack a small molecule binding pocket in its free form structure. The inhibition of PHF20L1 paves an alternative way to target DNMT1 degradation.

## Introduction

Post-translational modifications (PTMs) of proteins play versatile roles in the regulation of almost every aspect of cellular function. For instance, gene transcription is exquisitely regulated by lysine acetylation and methylation on histone tails [1, 2]. Recently, non-histone protein methylation has gained increasing interest in the regulation of other protein modifications, protein-protein interactions, protein stability, subcellular localization and promoter binding [3, 4]. Similar to histone PTMs, non-histone protein methylation is also regulated by modification enzymes to write and erase methylation markers, as well as effector proteins, such as Tudor domain-containing proteins (TDRDs), to recognize and bind to methyl-lysine/arginine residues [5, 6].

PHD finger protein 20-like protein 1 (PHF20L1) contains two tandem Tudor domains on its N-terminus to read methyl-lysines of multiple non-histone proteins. Recognition of methylated pRb is localized at the E2F target genes, which in turn recruits the MOF acetyltransferase complex [7]. Consequently, the integrity of a pRb-dependent G1-S-phase checkpoint is maintained by the interplay with PHF20L1. Recently, a novel function of non-histone protein methylation is to trigger the proteolysis of methylated proteins. In human ovarian teratocarcinoma PA-1 cells, PHF20L1 and the lysine demethylase LSD1 coordinate to protect SOX2 from methylation-dependent proteolysis [8, 9]. The first Tudor domain (Tudor1) of PHF20L1 is ascribed to read not only the mono-methylated lysine K4 of histone H3 and K20 of histone H4 but also mono-methylated K142 (K142^me1^) of non-histone protein DNA (cytosine-5) methyltransferase 1 (DNMT1) [10, 11]. During DNA replication, DNMT1 methylates the newly synthesized daughter strand to preserve the CpG DNA methylation patterns [12]. Residue K142 of DNMT1 is methylated by the methyltransferase SET7, and this modification in turn recruits L3MBTL3 and ubiquitin ligase CRL4 (DCAF5) for the proteasomal degradation of DNMT1 [13–15]. PHF20L1 competes with L3MBTL3 to antagonize DNMT1 degradation primarily in S phase [16].

DNMT1 is the major enzyme responsible for maintaining the epigenetic inheritance of the DNA cytosine methylation pattern and has gained tremendous therapeutic interest [17–19]. Two nucleoside-like drugs (azacitidine and decitabine) that trigger the proteasomal degradation of DNMT1 have been approved for the treatment of myeloid malignancies [20]. Subsequently, several nucleoside-like DNMT1 inhibitors have been designed as prodrugs, e.g., CP-4200 (an elaidic acid ester of azacitidine) and SGI-110 (a dinucleotide decitabine-p-deoxyguanosine), for better stability and lower toxicity [21, 22]. To enhance the specificity for DNMT1, non-nucleoside inhibitors are now in preclinical studies or clinical trials, which do not incorporate into genomic DNA and show lower cytotoxicity. Due to the regulatory roles of PHF20L1 in DNMT1 degradation, aberrations in PHF20L1 gene expression are highly correlated with various cancers, e.g., breast and ovarian cancers [23, 24]. PHF20L1 is the most commonly amplified TDRD gene in TCGA breast cancers, in which cell proliferation is suppressed by knockdown of PHF20L1 [25]. PHF20L1, especially its first Tudor domain, thus becomes an emerging therapeutic target.

Structural studies of Tudor domains demonstrate that three to five aromatic residues pile together to form an aromatic cage to accommodate the methylated lysine/arginine [26–29]. However, the PHF20 Tudor domain, as a homolog of PHF20L1, in its free form adopts another closed form conformation with the aromatic cage blocked by the flipped aromatic rings [30]. The molecular mechanism underlying the ligand recognition of PHF20L1 Tudor1 thus remains enigmatic. This also brings out an interesting question regarding the druggability of proteins lacking small molecule binding pockets in their free form. Here, we solved the crystal structure of the free form of PHF20L1 Tudor1, whose aromatic cage adopts a closed conformation similar to PHF20. NMR relaxation dispersion and molecular dynamic (MD) simulations suggest the pre-existence of another low-population conformation of the free form PHF20L1 Tudor1. The crystal structure of PHF20L1 Tudor1 in complex with K142^me1^ DNMT1 and the cosolvent MES reveals that the residues proximal to the aromatic cage are dramatically rearranged to accommodate the methyl-lysine. The same open form conformation of PHF20L1 Tudor1 was adopted in its crystal structure in complex with NMR fragment screening hit **1**. Mutations of key residues modulate the affinity for hit **1** by altering the structure and dynamics of the aromatic cage residues. These PHF20L1 mutants were further assessed by their cellular co-localization with DNMT1. Our work elucidates the conformational selection mechanism of PHF20L1 Tudor1 by which it recognizes both the natural substrate DNMT1 and small molecules. This study shall also have a profound effect on the druggability assessment of flexible targets, even though the free form structure lacks any binding sites for chemical probes.

## Results

### The PHF20L1 Tudor1 structure reveals a closed form aromatic cage

We first solved the free form crystal structure of PHF20L1 Tudor1 (Figure S1, Table S1), which reveals a canonical Tudor fold with four-stranded anti-parallel β sheets (β_1_-β_4_). Sequence alignment with previously published structures suggests that PHF20L1 Tudor1 is homologous other Tudor and MBT domains (Figure S2), which all belong to the Royal superfamily. These domains utilize three to five aromatic residues to form an aromatic cage to accommodate methyl-lysine. Although the aromatic planes of residues Y24, Y29 and Y54 of PHF20L1 are roughly perpendicular to that of F47, the side chain of W50 forms a π–π interaction with F47 and thus blocks the aromatic cage (Figure S1a and S1b). Such blocked conformation is similar to the solution structure (PDB code: 2EQM and 2JTF) of the free-form PHF20L1 Tudor1 (Figure S1c). This blocked aromatic cage is also observed in the PHF20 Tudor domain, which shares a 70 % sequence identity with PHF20L1 Tudor1. However, this static structure with the closed aromatic cage brings out an apparent paradox with respect to the fact that PHF20L1 Tudor1 recognizes several lysine-methylated substrates.

### Conformational plasticity of PHF20L1 Tudor1

Two possible mechanisms may unveil the enigma: first, PHF20L1 Tudor1 may have a preexisting low-population conformation to accommodate methyl-lysine (conformational selection [31–34]); or second, the lysine-methylated substrates may induce conformational changes to PHF20L1 Tudor1 (induced fit [35]). NMR relaxation dispersion experiments have proven to be fruitful for probing the preexisting minor states [36–38]. We hence first assigned the backbone chemical shifts of PHF20L1 using heteronuclear three-dimensional NMR spectroscopy. A total of 59 residues were unambiguously assigned except for Y24 and W50, which were missing signals in the ^1^H-^15^N heteronuclear single-quantum coherence (HSQC) spectra, and 5 prolines (Figure S3). The variable temperature NMR HSQC spectra demonstrate that the intensities of some aromatic cage residues drop more significantly than others (Figure S4), indicating that these residues may undergo conformational exchanges.

To probe the conformational plasticity, NMR relaxation dispersion (RD) experiments based on Carr– Purcell–Meiboom–Gill (CPMG) sequences were carried out for the ^15^N-labeled PHF20L1 Tudor1 (Figure 1a). RD has gained extensive application in studies of protein folding, enzyme catalysis, molecular recognition and ligand binding [38–45]. In cases where the residues are in interconversion on the millisecond time scale between the ground state and a low-population excited state, the RD profile (the effective transverse relaxation constant 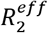) varies relative to the CPMG frequency, defined as the reciprocal of the delay between consecutive inversion pulses during the relaxation period. Conversely, the residues not undergoing conformational changes on this millisecond time scale show flattened RD profiles. It is interesting to observe that residues Y29, F47 and Y54 in the closed aromatic cage are in dynamic exchange with another state, as suggested by the global fitting of these RD profiles. Residue W50 is missing in the HSQC spectra, probably due to the exchange-induced line broadening. Conversely, residues distal to the aromatic cage show no conformational changes, at least on the millisecond time scale of the RD experiments.

**Figure 1.**
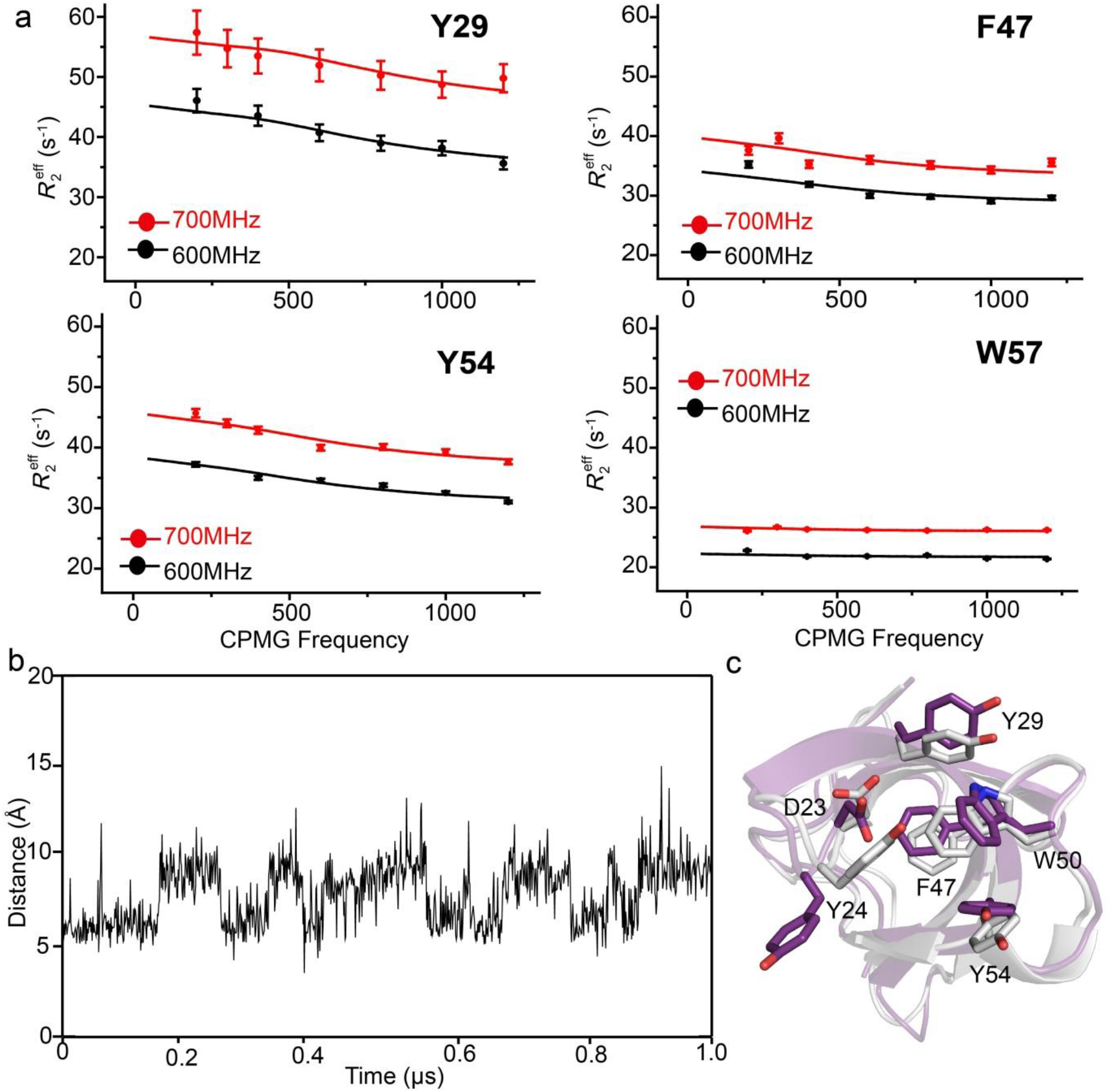
Conformational exchange of the apo-PHF20L1 Tudor1. **a**) Representative residues from the relaxation dispersion profiles. Solid lines denote global fitting using the two-state model. Error bar represents 2σ estimated from the signal to noise ratio of each NMR peak. **b**) Aromatic cage residues of PHF20L1 Tudor1 interconvert between different states, as characterized by the distance between the Y24 C^γ^ atom and the W50 C^η2^ atom. The conformations of these two residues are remarkably changed during molecular dynamics simulations. **c**) Conformational changes of the aromatic cage residues of PHF20L1 Tudor1 during molecular dynamics simulations. Interconversion between the closed crystal structure (gray) and the open representative structures from 1us MD simulation (purple) of the closed form.

To further explore the conformational exchange of PHF20L1 Tudor1, we carried out a 1 μs all-atom MD simulation starting from the free form crystal structure. A remarkable conformational rearrangement of the aromatic cage residues e.g., residues Y24 and W50, was observed. We hence defined the Y24-W50 distance as an indicator of the conformational exchange in MD frames (Figure 1b). Clearly, the free form PHF20L1 Tudor1 switches between two states, one of which is the native closed form, and another is an open form (Figure 1c) that can potentially be deployed to recognize the mono-methylated substrate.

### PHF20L1 Tudor1 recognizes K142 mono-methylated DNMT1

We hence determined the binding between PHF20L1 Tudor1 and the natural substrate K142^me1^ DNMT1 using NMR chemical shift perturbations (CSPs). As expected, the titration of the K142^me1^ DNMT1 peptide induced significant chemical shift perturbations of ^15^N-labeled PHF20L1 (Figure 2a), especially for residues in the aromatic cage (Figure 2b). The binding affinity (*K*_D_ = 0.67 mM) was derived from the dose-dependent CSPs, which was consistent with that determined from isothermal titration calorimetry (Figure 2c). We then determined the crystal structure of PHF20L1 Tudor1 in complex with the K142^me1^ DNMT1 peptide, diffracted at a resolution of 1.90 Å (Table S1). The electron densities of the residues from R140 to G145 of DNMT1 can clearly be observed in the complex structure (Figure 2d). The methylated K142 protrudes into the aromatic cage surrounded by residues Y24, Y29, F47, W50 and Y54 (Figure 2e), and the N^ε^ atom of K142 also forms a direct hydrogen bond with D23 and a water-bridged hydrogen bond with E56 (Figure 2f). In addition, R140 of DNMT1 forms two hydrogen bonds with R49 and Y54 of PHF20L1. Although the complex structure in general superimposed well with respect to free form PHF20L1 Tudor1 at a r.m.s.d. of 0.43 Å for C_α_ atoms, residues from the C-terminus of β_1_ to the N-terminus of β_2_ were dramatically rearranged. For instance, residues D23, Y24 and Y29 were remarkably rearranged relative to the free form PHF20L1 Tudor1 structure (Figure 2g). Such a conformational rearrangement enables preferential binding to DNMT1 K142^me1^, e.g., the carboxyl group of D23 was displaced approximately 4.0 Å to form a direct hydrogen bond with the N^ε^ atom of DNMT1 K142, while the phenol groups of Y24 and Y29 were displaced 7.9 and 5.8 Å away to form one hydrogen bond with DNMT1 K142 and one water-mediated hydrogen bond with D23, respectively (Figure 2h). Such conformational changes facilitate W50 to turn over approximately 74°outward to stack with K142 and R140 of DNMT1 in a sandwich-like mode and open the aromatic cage to accommodate K142^me1^ DNMT1.

**Figure 2.**
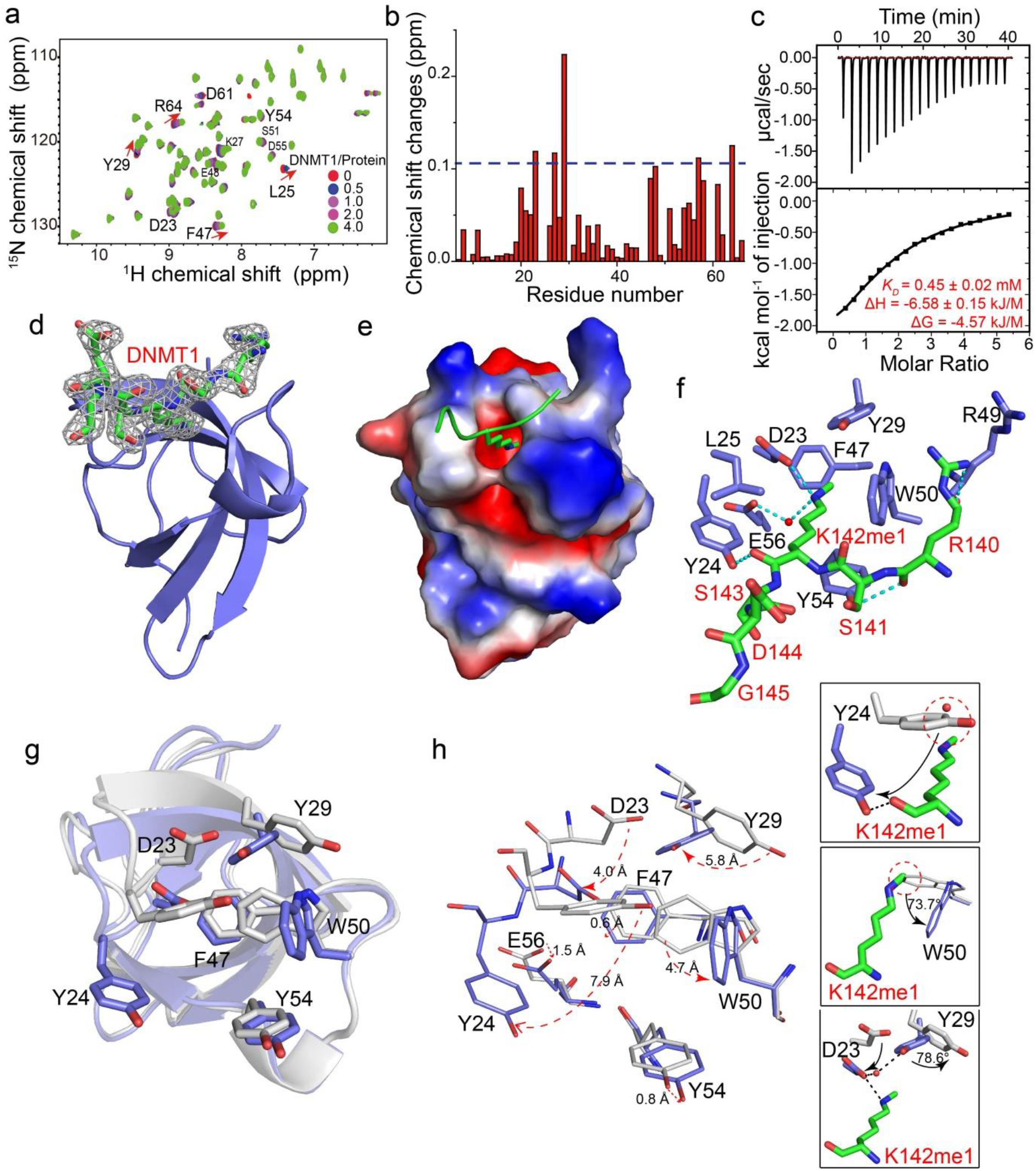
PHF20L1 Tudor1 recognizes the natural peptide substrate K142^me1^ DNMT1 through conformational rearrangement of aromatic cage residues. **a**) Chemical shift perturbations of ^15^N-labeled PHF20L1 Tudor1 induced by the K142^me1^ DNMT1 peptide. **b**) Residue-by-residue chemical shift changes of PHF20L1 Tudor1 at a peptide/protein molar ratio of 4:1. The dashed line denotes one standard deviation above the average chemical shift changes of the residues, excluding Y29. **c**) The binding affinity of the K142me1 DNMT1 peptide was determined by isothermal titration calorimetry. **d**) Cartoon representation of PHF20L1 Tudor1 in complex with K142^me1^ DNMT1 (carbon atoms in green). The 2Fo-Fc electron-density map of the K142^me1^ DNMT1 peptide was contoured at 1.0 σ (gray). **e**) The electrostatic surface representation of PHF20L1 Tudor1, in which the mono-methylated K142 is depicted as sticks. **f**) The detailed interactions between PHF20L1 Tudor1 (slate) and the K142^me1^ DNMT1 peptide (green). Hydrogen bonds are denoted by dashed lines. **g**) Superimposition of the crystal structures of the free (gray) and bound (slate) forms of PHF20L1 Tudor1. **h**) Conformational rearrangement of the aromatic cage residues of the DNMT1-bound PHF20L1 Tudor1 relative to those of the free form. Arrows denote the spatial displacements (values annotated) of the aromatic cage residues. The aromatic ring flips and displacements of residues such as D23, Y24, Y29 and W50 allow the protrusion of methyl-K142 of DNMT1.

The open and closed form populations were thereafter estimated from MD simulations using the distance between the Y24 C^γ^ atom and W50 C^η2^ atom in the complex crystal structure (10.2 Å) as a threshold. This suggests that the open form has a low population of approximately 6 % (Figure 1b), which is in good agreement with the population of 6.8 ± 0.04 % (fitting error) estimated from the global fitting of the relaxation dispersion profiles of the apo-PHF20L1 Tudor1. The low-population open form structure from the MD simulations is also similar to the complex crystal structure of PHF20L1 Tudor1 (Figure 1c). These data underpin that PHF20L1 Tudor1 utilizes the preexisting low-population open form conformation to recognize the natural substrate K142^me1^ DNMT1.

### MES binds to the open form aromatic cage of PHF20L1 Tudor1

During our structural studies of PHF20L1 Tudor1, we fortuitously solved the crystal structure of PHF20L1 Tudor1 in complex with the solvent molecule 2-(N-morpholino)ethanesulfonic acid (MES), diffracted at a resolution of 1.60 Å (Figure S5a, Table S1). MES binds to the open form aromatic cage of PHF20L1 Tudor (Figure S5b), which adopts a similar conformation relative to that of the PHF20L1-DNMT1 complex (Figure S5c), at a r.m.s.d. of 0.39 Å for C_α_ atoms. MES induces detectable CSPs of PHF20L1 Tudor1 at a very high molar ratio of 1000:1 but almost no changes are observed at the ratio of 24:1 (Figure S6). Although PHF20L1 binds MES with a very weak affinity, this result sheds light on the potential druggability of the low-population open form aromatic cage of PHF20L1 Tudor1.

### NMR fragment-based screening against PHF20L1 Tudor1

We hence carried out fragment-based screening against PHF20L1 Tudor1. Although the saturation transfer difference and WaterLOGSY spectra have been extensively applied to identify small molecule hits, the low molecular weight of PHF20L1 Tudor1 may lower the sensitivities of these ligand-observed spectra. Instead, the protein-observed CSPs were utilized to uncover hits from 89 cocktails with 10 compounds each to improve screening throughput. Each component of the cocktails inducing significant CSPs (Figure S7) was then titrated to ^15^N-labeled PHF20L1 Tudor1 for hit deconvolution. A total of three hits were thus identified to bind to the aromatic cage. For instance, hit **1** induces dose-dependent CSPs (Figure 3a), particularly for residues proximal to the aromatic cage of PHF20L1 (Figure 3b). Although a weak binding affinity of 0.31 mM was determined from the dose-dependent CSPs (Figure 3c), the remarkably large CSP of 0.65 ppm for residue L25 suggests conformational rearrangement of the aromatic cage of PHF20L1 Tudor1.

**Figure 3.**
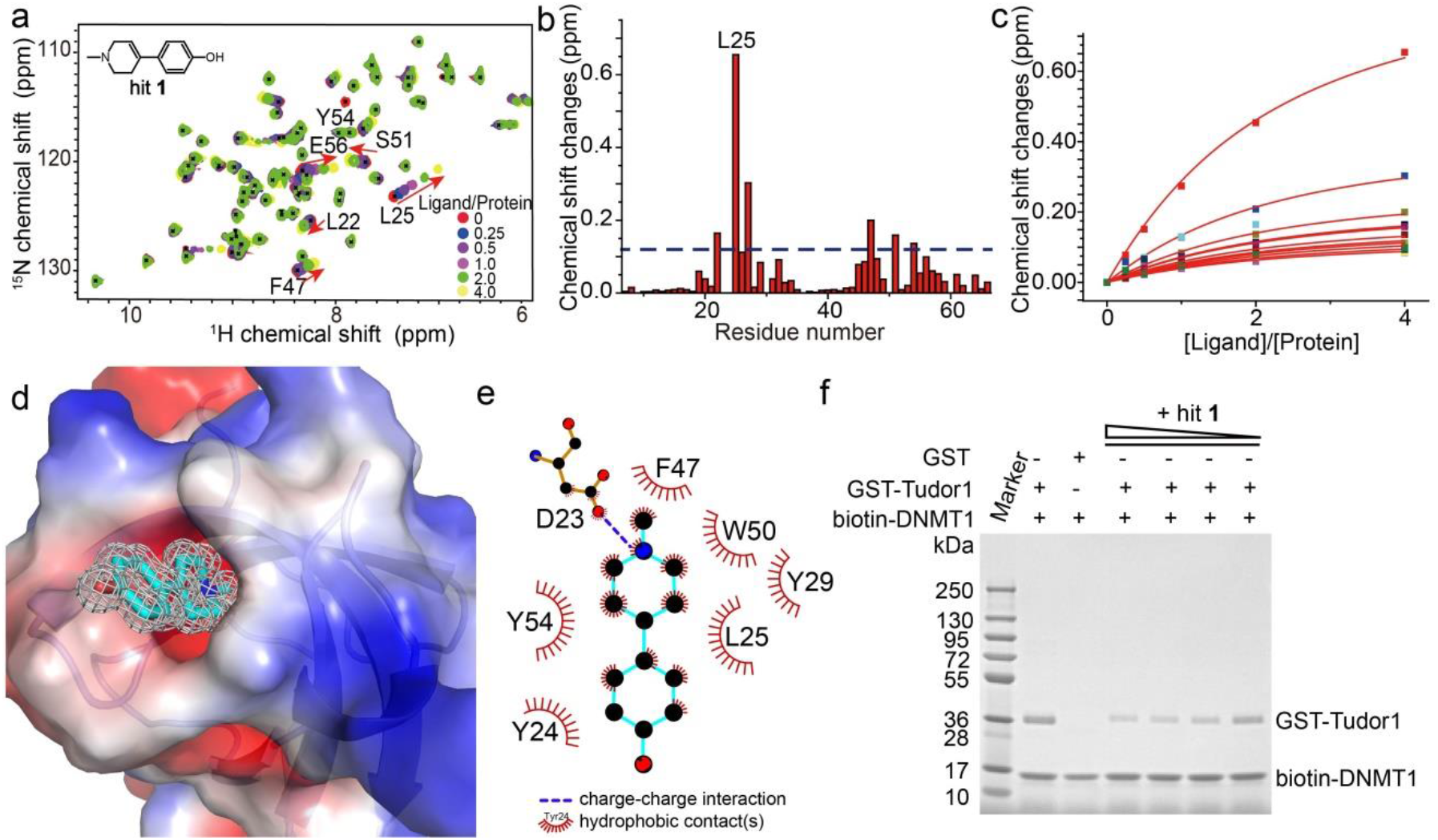
Fragment-based screening hit **1** binds to the open form aromatic cage of PHF20L1 Tudor1. **a**) The chemical shift perturbations of ^15^N-labeled PHF20L1 Tudor1 upon titration of hit **1** at various ligand/protein molar ratios. **b**) Residue-by-residue chemical shift changes of PHF20L1 Tudor1 domain at a ligand/protein molar ratio of 4:1. The dashed line denotes one standard deviation above the average chemical shift changes of residues, excluding L25. **c**) The binding affinity of hit **1** is best fitted from the dose-dependent chemical shift changes of PHF20L1 Tudor1. **d**) The electrostatic surface representation of PHF20L1 Tudor1 in complex with hit **1** (carbon atoms in cyan). The 2Fo-Fc electron-density map of hit **1** was contoured at 1.0 σ (gray). **e**) Detailed interactions between hit **1** and PHF20L1 Tudor1 analyzed using the LIGPLOT program. **f**) The pull-down assay of the biotin-labeled K142^me1^DNMT1-peptide (annotated as biotin-DNMT1) against the GST-PHF20L1 Tudor1 (abbreviated as GST-Tudor1) upon the titration of hit **1**, at the hit **1**: GST-Tudor1 molar ratio of 100, 50, 25, and 12.5, respectively.

The crystal structure of PHF20L1 Tudor1 in complex with hit **1** with clear electron density reveals that this small molecule also protrudes into the open form aromatic cage (Figures 3d, S8a, Table S1). More importantly, the methylated nitrogen atom of hit **1** mimics the N^ε^ atom of K142^me1^ DNMT1 (Figure S8b), and both were found deviating approximately 2.8 Å away from the carboxyl oxygen of D23 of PHF20L1 Tudor1 (Figure 3e). Such a strong electrostatic interaction explains the much weaker affinity of MES, as its nitrogen atom was found deviating 3.5 Å away. Once again, Y24 and W50 are the essential residues that regulate the interconversion between the open and closed states of the aromatic cage of PHF20L1 Tudor1. The pull-down assay of the biotin-labeled K142^me1^ DNMT1 peptide and PHF20L1 Tudor1 demonstrated that the hit **1** blocks this PHF20L1-DNMT1 interaction in a dose-dependent manner (Figure 3f), indicating the druggability of PHF20L1 Tudor1.

### Molecular mechanism of the open-to-closed conformational exchange of the aromatic cage

To further validate the regulatory role of these aromatic cage residues in the ligand recognition of PHF20L1 Tudor1, we carried out mutagenesis studies of key residues (Table S2). Considering the intermolecular cation-π and hydrophobic interactions, it is not surprising that W50A and Y54A mutations abolish the binding to hit **1**. However, the Y24A or Y24L mutants have an almost identical binding affinity for hit **1** even though weakened cation-π and hydrophobic interactions are expected. On the other hand, although Y29W mutagenesis only slightly weakens the binding to hit **1**, the Y24W mutant binds at a much weaker affinity of 1.5 mM. Interestingly, the R49G and R49W mutant has about 1.7 and 1.4 fold enhancement of the ligand-binding capacity, respectively, although residue R49 is not involved in ligand recognition as revealed in the crystal structures of PHF20L1 Tudor1 in complex with hit **1**(Figure 3e). That is to say, the static free and bound form crystal structures of PHF20L1 cannot explain the structure-activity relationships, as mutagenesis may alter the topology of the aromatic cage and/or the conformational dynamics.

We hence determined the free-form crystal structure of PHF20L1 Tudor1 Y24L and Y24W/Y29W mutants (Figure 4a, 4b, and Table S3). In the wild-type free form structure of PHF20L1 Tudor1, the side chain phenol group of Y24 closely interacts with the aromatic group of W50, with a minimum atom-to-atom distance of 3.5 Å (Figure S1a). In the Y24L mutant structure, the less bulky and more flexible side chain of L24 allows W50 to adopt an open form conformation (Figure 4a) similar to the conformations observed in the PHF20L1 complexes. That is, despite a weakened cation-π interaction for the Y24L mutant, preferential switching to the open form conformation compensates for the ligand binding capability. However, the aromatic cage in the Y24W/Y29W mutant is blocked by the T-stacking interaction between Y24W and W50 (Figure 4b).

**Figure 4.**
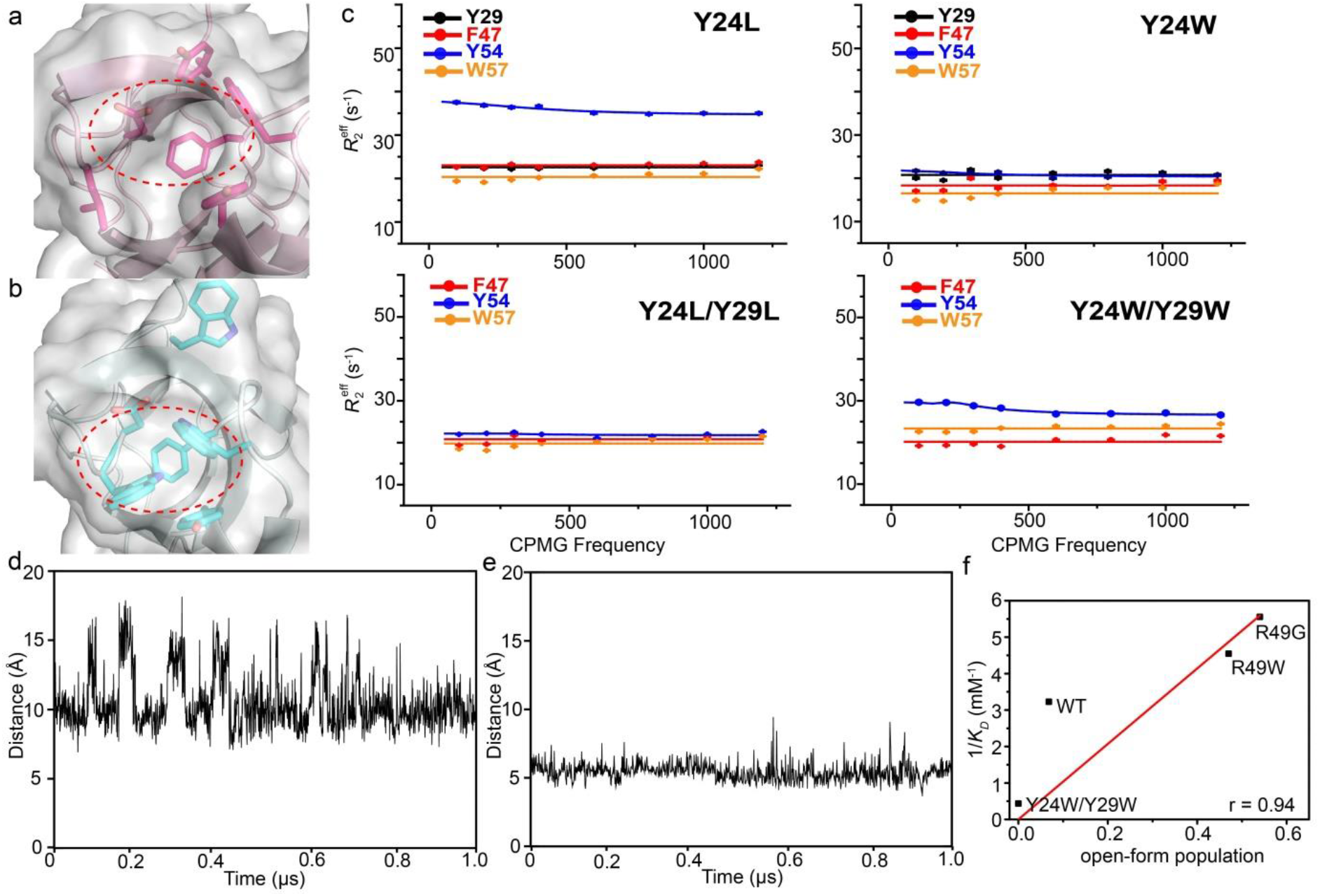
Mutations of key residues in the aromatic cage alter both the free form structure and open-to-closed conformational exchanges. **a**) The open form aromatic cage (highlighted in the red oval) of the Y24L mutant of PHF20L1 Tudor1. **b**) The crystal structure of the Y24W/Y29W mutant of PHF20L1 Tudor1, in which the aromatic cage is blocked by T-stacking of Y24W and W50. **c**) Relaxation dispersion profiles for typical residues in various mutants as annotated. Error bar of 2σ was estimated from the NMR signal to noise ratio of each residues. **d**) and **e**) MD simulations of the Y24L and Y24W/Y29W mutants. **f**) The association constant (1/*K*_D_) correlates with the open-form population. Annotated is the Pearson’s coefficient when fitting to a directly proportional function.

The aforementioned analysis indicates that there is a correlation between ligand binding capacity and conformational distribution, which can be quantitatively described assuming a two-step binding equilibrium (Eq. S2-S4). It suggests that the apparent binding affinity can be regulated by the population of the open state reciprocally. The conformational dynamics of these mutants were then analyzed by NMR relaxation dispersion experiments. Interestingly, the effective transverse R2 rates of the aromatic cage residues of these mutants were significantly reduced relative to those of the wild-type protein, e.g., the effective R2 of Y54 in the low CPMG frequency drops from approximately 45 Hz in the wild-type to 38, 23, 17, and 27 Hz in the Y24L, Y24W, Y24L/Y29L and Y24W/Y29W mutants, respectively. Flattened relaxation dispersion profiles were also observed for these free-form mutants (Figure 4c), which suggest that the conformation exchange either is not on the millisecond time scale or is suppressed by these mutations. Excess K142me1 DNMT1 peptide also flattened the relaxation dispersion profile (Figure S9), as in such condition the population of the open state was remarkably reduced.

Hence, we performed MD simulations with the Y24L and Y24W/Y29W mutants of PHF20L1 Tudor1 starting from the mutant crystal structures (Figure 4d, 4e). Using the same distance threshold defined previously, we estimated that the open form of the Y24L mutant has a higher population (67%) than the wild-type one (6%). That is, Y24L mutagenesis tends to switch towards the open form conformation. This switch compensates for the binding energy loss due to the Y24L mutation; hence, similar ligand-binding affinities were observed for the wild-type and the Y24L mutant (Table S2). Conversely, the Y24W/Y29W mutagenesis completely trapped the aromatic cage in a closed form in our simulations; thus, a weak ligand-binding capability was observed. Interestingly, R49G and R49W mutagenesis shift the open-form population to 54% and 47%, respectively. This offers a way to investigate the contribution of conformational distribution separately, as R49 is not directly interacting with hit **1**. For this reason, the fragment screening hit is a better probe of conformational dynamics than the peptide, as the flanking residue R140 of the DNMT1 peptide are in close contact with R49 of PHF20L1 Tudor1. The monotonous increment of association constant (1/*K*_D_) relative to the open-form population (Figure 4f), as predicted by the conformational selection model (Eq. S4).

Principal component analysis (PCA) was carried out to extract the contact patterns of the two mutants during MD simulations (Figure S10) [46]. PCA was performed using the heavy atoms of residues D23, Y24, L25, Y29, F47, W50 and Y54 to yield PCA modes describing representative motions of closed state. Projects the trajectory of closed state (1004 structures in total) onto the 2D essential subspace defined by the first two PCs, and the two crystal structures were also projected onto the plane (Figure S10a). These two PCA modes contribute ~42% and ~21% of the total fluctuation in the WT, respectively. The projection of the closed crystal structure (red) and the open crystal structure (blue) are (16.6, −21.0) and (27.3, −23.0), respectively (Figure S10a). There are mainly open states on the right side of the blue dot with cutoff 10.2Å of Y24-W50, and mainly closed states on the left side of the blue dot. That is to say, there is closed-open transition during 1 μs MD simulation. We further performed contact-based PCA, in order to detect key contacts that control the closed-open transition. The elements of the normalized eigenvector of the first PC of the closed state (black), the Y24L mutant (red) and the Y24W/Y29W mutant (blue) (Figure S10b). In the closed state, the contacts between Y24-Y54 and Y24-W50 mainly control the close-open transition of the protein, whereas other contacts make little contribution to the conformational transition. The Y24L mutant is prone to take the open state than the closed state, the weights of the L24-Y54 and L24-W50 contacts decrease, but there are quite some other contacts, such as L25-Y54, D23-Y54, Y29-Y54, F47-Y54 and W50-Y54, make much contribution to this mode. This may promote to adopt an open form conformation for Y24L. There is no close-open transition in the Y24W/Y29W mutant. Therefore W24-Y54 and W24-W50 have little contribution to this mode. Instead contacts, such as L25-W54 and D23-W54, have a larger contribution than those in the WT, but they are not essential in conformational transition.

### PHF20L1 Tudor1 recognizes DNMT1 in cells

To validate whether PHF20L1 Tudor1 recognizes DNMT1 in a cellular context, we first carried out immunofluorescence experiments for various PHF20L1 mutants in Hela cells (Figures 5a and S11a). The wild-type PHF20L1 and its DNMT1-binding mutants showed dispersed distributions in the nucleus and colocalized with DNMT1, while the mutants lost their DNMT1 binding capability and form the isolated foci. The co-localization of wild type or mutant PHF20L1 with DNMT1 was statistically described by the spatial quantification of green and red fluorescence along a path across the nucleus (Figure 5b). The cellular co-localization study agrees well with the binding capability of these PHSF20L1 mutants (Table S2). Hence, the *in vitro* mutagenesis studies were well recapitulated in the cellular assays.

**Figure 5.**
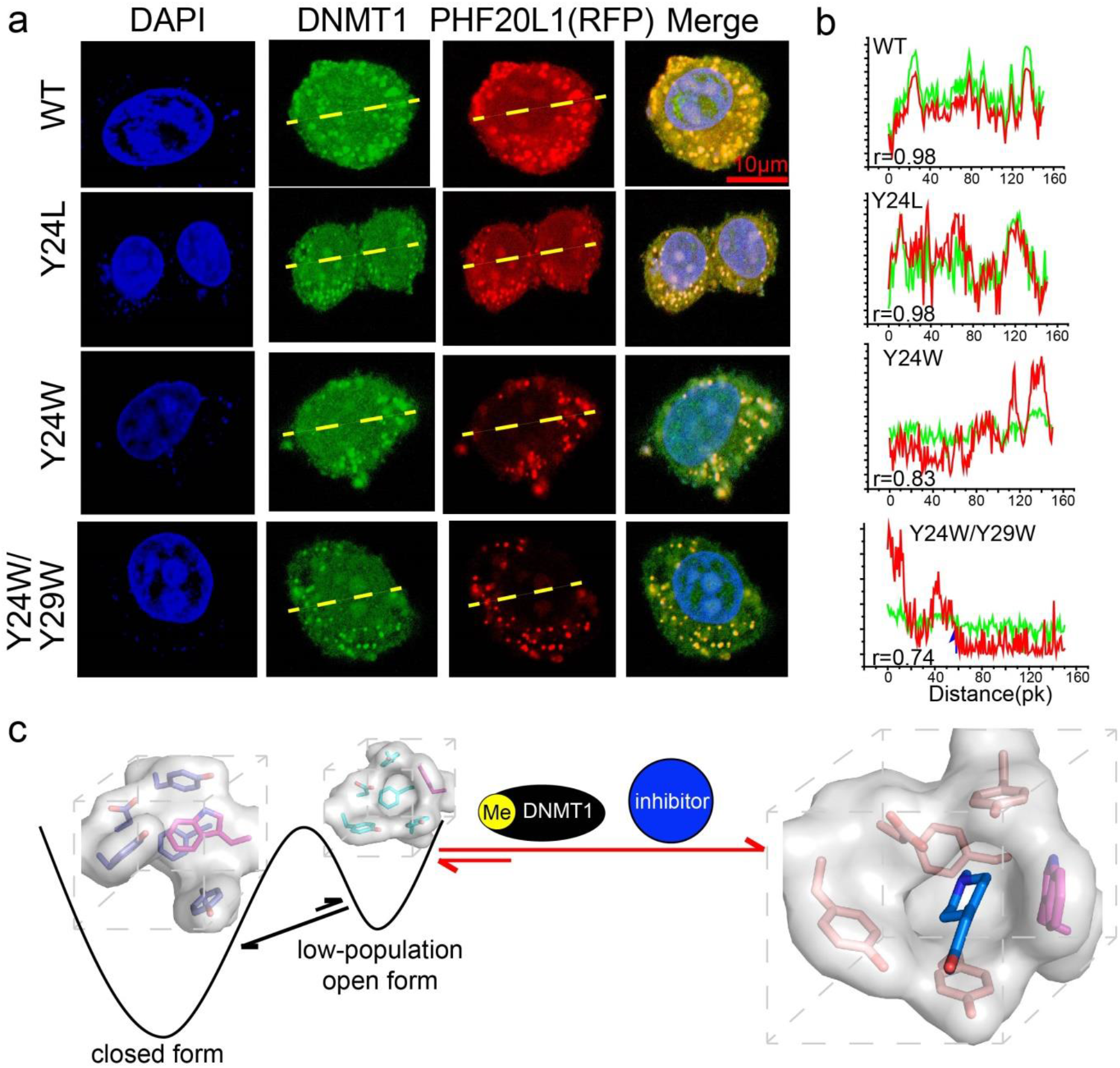
**a**) Immunofluorescence assay of endogenous DNMT1 (stained in green by antibodies) with ectopically overexpressed RFP-fused PHF20L1 or its mutants in HeLa cells. **b**) Spatial quantification was performed along a path across the nucleus (a white line in the merged images in the left), along which Green and red fluorescence was quantified. Annotated was the statistical Person’s coefficient to describe the co-localization of wild type or mutant PHF20L1 with DNMT1. **c**) Conformational selection model of PHF20L1 Tudor1 for its ligand recognition.

## Discussion

By combining X-ray crystallography, NMR and MD simulations, we uncovered the conformational selection mechanism underlying the ligand recognition of PHF20L1 Tudor1. Previous studies have shown that small molecule binding can lead to dramatic conformational differences that typically reflect dynamics and selection of pre-existing conformational states [47–51]. Since the population of the open and closed forms can be shifted by temperature, mutagenesis, cofactors, etc., this entitles PHF20L1 an extra level to mediate ligand binding. PHF20L1 binds not only to methylated DNMT1 but also to mono-methylated pRb, which is essential for the pRb-dependent G1–S-phase checkpoint [7]. This indicates the interplay between conformational exchanges and the on-off switch of biological functions in a spatiotemporal manner. On the other hand, the conformational selection mechanism entitles the discovery of small molecule hits against the seemingly “undruggable” PHF20L1 Tudor1. Our results emphasize the importance of protein dynamics on the druggability assessment of a potential target (Figure 5c), even though it may present an “undruggable” conformation in the free form. Although not at the lowest energy level, the minor conformation exists at a low but detectable population. As the same open form conformation of PHF20L1 has been selected to recognize both the natural substrate and synthesized compounds, following hit-to-lead evolution shall benefit from this low-population open form conformation.

## Acknowledgments

We thank the staff at the BL17B/BL18U1/BL19U1 beamline of the National Center for Protein Sciences Shanghai (NCPSS) at the Shanghai Synchrotron Radiation Facility. Part of our NMR work was performed at the National Center for Protein Sciences Shanghai and High Magnetic Field Laboratory of the Chinese Academy of Sciences. We thank the support of the Supercomputing Center of USTC and the Bioinformatics Center of the School of Life Sciences at USTC. This study was financially supported by the Ministry of Science and Technology of China (2016YFA0500700 and 2019YFA0508400), the Natural Science Foundation of China (21874123, 21573205, 21807095), Hefei National Science Center Pilot Project Funds, Fundamental Research Funds for the Central Universities (WK2060190086), the New Concept Medical Research Fund of USTC (to YP), China Postdoctoral Science Foundation (No. 2019M662182).

## Database

Structural data are available in the PDB under accession number 6L0X, 6L1F, 6L10, 6L1P, 6L1C, 6L1I.

## References

1. Strahl, B.D. and C.D. Allis, The language of covalent histone modifications. Nature, 2000. 403(6765): p. 41–45.

2. Huang, H., et al., Quantitative Proteomic Analysis of Histone Modifications. Chemical Reviews, 2015. 115(6): p. 2376–2418.

3. Hamamoto, R., V. Saloura, and Y. Nakamura, Critical roles of non-histone protein lysine methylation in human tumorigenesis. Nature Reviews Cancer, 2015. 15(2): p. 110–124.

4. Biggar, K.K. and S.S.C. Li, Non-histone protein methylation as a regulator of cellular signalling and function. Nature Reviews Molecular Cell Biology, 2015. 16(1): p. 5–17.

5. Musselman, C.A., et al., Perceiving the epigenetic landscape through histone readers. Nature Structural & Molecular Biology, 2012. 19(12): p. 1218–1227.

6. Lu, R. and G.G. Wang, Tudor: a versatile family of histone methylation ‘readers’. Trends In Biochemical Sciences, 2013. 38(11): p. 546–555.

7. Carr, S.M., et al., Tudor-domain protein PHF20L1 reads lysine methylated retinoblastoma tumour suppressor protein. Cell Death And Differentiation, 2017. 24(12): p. 2139–2149.

8. Wang, Q.Q., et al., PHF20L1 antagonizes SOX2 proteolysis triggered by the MLL1/WDR5 complexes. Laboratory Investigation, 2018. 98(12): p. 1627–1641.

9. Zhang, C.X., et al., LSD1 demethylase and the methyl-binding protein PHF20L1 prevent SET7 methyltransferase-dependent proteolysis of the stem-cell protein SOX2. Journal Of Biological Chemistry, 2018. 293(10): p. 3663–3674.

10. Kim, J., et al., Tudor, MBT and chromo domains gauge the degree of lysine methylation. Embo Reports, 2006. 7(4): p. 397–403.

11. Esteve, P.O., et al., Methyllysine Reader Plant Homeodomain (PHD) Finger Protein 20-like 1 (PHF20L1) Antagonizes DNA (Cytosine-5) Methyltransferase 1 (DNMT1) Proteasomal Degradation. Journal Of Biological Chemistry, 2014. 289(12): p. 8277–8287.

12. Esteve, P.O., et al., Direct interaction between DNMT1 and G9a coordinates DNA and histone methylation during replication. Genes & Development, 2006. 20(22): p. 3089–3103.

13. Esteve, P.O., et al., Regulation of DNMT1 stability through SET7-mediated lysine methylation in mammalian cells. Proceedings Of the National Academy Of Sciences Of the United States Of America, 2009. 106(13): p. 5076–5081.

14. Leng, F., et al., Methylated DNMT1 and E2F1 are targeted for proteolysis by L3MBTL3 and CRL4(DCAF5) ubiquitin ligase. Nature Communications, 2018. 9.

15. Esteve, P.O., et al., A methylation and phosphorylation switch between an adjacent lysine and serine determines human DNMT1 stability. Nature Structural & Molecular Biology, 2011. 18(1): p. 42-+.

16. Zhang, C.X., et al., Proteolysis of methylated SOX2 protein is regulated by L3MBTL3 and CRL4(DCAF5) ubiquitin ligase. Journal Of Biological Chemistry, 2019. 294(2): p. 476–489.

17. Sharif, J., et al., The SRA protein Np95 mediates epigenetic inheritance by recruiting Dnmt1 to methylated DNA. Nature, 2007. 450(7171): p. 908–U25.

18. Cortez, C.C. and P.A. Jones, Chromatin, cancer and drug therapies. Mutation Research-Fundamental And Molecular Mechanisms Of Mutagenesis, 2008. 647(1-2): p. 44–51.

19. Pradhan, S., et al., Recombinant human DNA (cytosine-5) methyltransferase I. Expression, purification, and comparison of de novo and maintenance methylation. Journal Of Biological Chemistry, 1999. 274(46): p. 33002–33010.

20. Gros, C., et al., DNA methylation inhibitors in cancer: Recent and future approaches. Biochimie, 2012. 94(11): p. 2280–2296.

21. Brueckner, B., et al., Delivery of 5-Azacytidine to Human Cancer Cells by Elaidic Acid Esterification Increases Therapeutic Drug Efficacy. Molecular Cancer Therapeutics, 2010. 9(5): p. 1256–1264.

22. Chuang, J.C., et al., S110, a 5-Aza-2 ‘-Deoxycytidine-Containing Dinucleotide, Is an Effective DNA Methylation Inhibitor In vivo and Can Reduce Tumor Growth. Molecular Cancer Therapeutics, 2010. 9(5): p. 1443–1450.

23. Wrzeszczynski, K.O., et al., Identification of Tumor Suppressors and Oncogenes from Genomic and Epigenetic Features in Ovarian Cancer. Plos One, 2011. 6(12).

24. Natrajan, R., et al., An integrative genomic and transcriptomic analysis reveals molecular pathways and networks regulated by copy number aberrations in basal-like, HER2 and luminal cancers. Breast Cancer Research And Treatment, 2010. 121(3): p. 575–589.

25. Jiang, Y.Y., et al., An integrated genomic analysis of Tudor domain-containing proteins identifies PHD finger protein 20-like 1 (PHF20L1) as a candidate oncogene in breast cancer. Molecular Oncology, 2016. 10(2): p. 292–302.

26. Taverna, S.D., et al., How chromatin-binding modules interpret histone modifications: lessons from professional pocket pickers. Nature Structural & Molecular Biology, 2007. 14(11): p. 1025–1040.

27. Dai, Y., et al., Structural basis for recognition of 53BP1 tandem Tudor domain by TIRR. Nat Commun, 2018. 9(1): p. 2123.

28. Botuyan, M.V., et al., Structural basis for the methylation state-specific recognition of histone H4-K20 by 53BP1 and Crb2 in DNA repair. Cell, 2006. 127(7): p. 1361–73.

29. Liu, J., et al., Structural plasticity of the TDRD3 Tudor domain probed by a fragment screening hit. FEBS J, 2018. 285(11): p. 2091–2103.

30. Cui, G.F., et al., PHF20 is an effector protein of p53 double lysine methylation that stabilizes and activates p53. Nature Structural & Molecular Biology, 2012. 19(9): p. 916–924.

31. Ma, B.Y., et al., Folding funnels and binding mechanisms. Protein Engineering, 1999. 12(9): p. 713–720.

32. Tsai, C.J., et al., Folding funnels, binding funnels, and protein function. Protein Science, 1999. 8(6): p. 1181–1190.

33. Tsai, C.J., B.Y. Ma, and R. Nussinov, Folding and binding cascades: Shifts in energy landscapes. Proceedings Of the National Academy Of Sciences Of the United States Of America, 1999. 96(18): p. 9970–9972.

34. Boehr, D.D., R. Nussinov, and P.E. Wright, The role of dynamic conformational ensembles in biomolecular recognition (vol 5, pg 789, 2009). Nature Chemical Biology, 2009. 5(12): p. 954–954.

35. Koshland, D.E., Application Of a Theory Of Enzyme Specificity To Protein Synthesis. Proceedings Of the National Academy Of Sciences Of the United States Of America, 1958. 44(2): p. 98–104.

36. Li, P.L., et al., Internal dynamics control activation and activity of the autoinhibited Vav DH domain. Nature Structural & Molecular Biology, 2008. 15(6): p. 613–618.

37. Yao, X.L., M.K. Rosen, and K.H. Gardner, Estimation of the available free energy in a LOV2-J alpha photoswitch. Nature Chemical Biology, 2008. 4(8): p. 491–497.

38. Boehr, D.D., et al., The dynamic energy landscape of dihydrofolate reductase catalysis. Science, 2006. 313(5793): p. 1638–1642.

39. Lange, O.F., et al., Recognition dynamics up to microseconds revealed from an RDC-derived ubiquitin ensemble in solution. Science, 2008. 320(5882): p. 1471–1475.

40. Baldwin, A.J. and L.E. Kay, NMR spectroscopy brings invisible protein states into focus. Nature Chemical Biology, 2009. 5(11): p. 808–814.

41. Chao, F.A., et al., Probing the Broad Time Scale and Heterogeneous Conformational Dynamics in the Catalytic Core of the Arf-GAP ASAP1 via Methyl Adiabatic Relaxation Dispersion. Journal Of the American Chemical Society, 2019. 141(30): p. 11881–11891.

42. Yuwen, T.R., et al., A Methyl-TROSY-Based H-1 Relaxation Dispersion Experiment for Studies of Conformational Exchange in High Molecular Weight Proteins. Angewandte Chemie-International Edition, 2019. 58(19): p. 6250–6254.

43. Alderson, T.R., et al., Local unfolding of the HSP27 monomer regulates chaperone activity. Nature Communications, 2019. 10.

44. Zhang, Q., et al., Visualizing spatially correlated dynamics that directs RNA conformational transitions. Nature, 2007. 450(7173): p. 1263–U14.

45. Henzler-Wildman, K.A., et al., Intrinsic motions along an enzymatic reaction trajectory. Nature, 2007. 450(7171): p. 838–U13.

46. Amadei, A., A.B.M. Linssen, and H.J.C. Berendsen, Essential Dynamics Of Proteins. Proteins-Structure Function And Genetics, 1993. 17(4): p. 412–425.

47. Wylie, A.A., et al., The allosteric inhibitor ABL001 enables dual targeting of BCR-ABL1. Nature, 2017. 543(7647): p. 733–737.

48. Jagtap, P.K.A., et al., Corrigendum: Selective Inhibitors of FKBP51 Employ Conformational Selection of Dynamic Invisible States. Angew Chem Int Ed Engl, 2019. 58(39): p. 13619.

49. Gaali, S., et al., Selective inhibitors of the FK506-binding protein 51 by induced fit. Nat Chem Biol, 2015. 11(1): p. 33–7.

50. Kunze, J., et al., Targeting dynamic pockets of HIV-1 protease by structure-based computational screening for allosteric inhibitors. J Chem Inf Model, 2014. 54(3): p. 987–91.

51. Kahraman, A., et al., Shape variation in protein binding pockets and their ligands. J Mol Biol, 2007. 368(1): p. 283–301.

