## Supplemental Information for "Conformational Selection in Ligand Recognition by the First Tudor Domain of PHF20L1"

### MATERIALS AND METHODS

#### Preparation of proteins and peptides

DNA encoding human PHF20L1 Tudor1 (residues 5-70) was amplified and incorporated into a modified pGEX-4T-1 (Novagen) plasmid, in which the thrombin protease site was substituted for a tobacco etch virus (TEV) cleavage site. All mutants of PHF20L1 were generated using a MutanBEST kit (Takara) and confirmed by DNA sequencing. The proteins were expressed in *Escherichia coli* Rosetta (DE3) cells (Novagen) cultured in LB medium or in $^{15}N/^{13}C$-enriched SV40 medium at 37 ℃ till $OD_{600}$ reaches about 1.2, then shifted to 16 ℃ and induced with 0.2 mM isopropyl-β-D-thiogalactopyranoside (IPTG) overnight. Bacterial pellets were resuspended in buffer A (20 mM Tris-HCl, 1 M NaCl, pH 8.0) and lysed by sonication on ice. The fusion proteins were purified on glutathione-Sepharose beads (GE Healthcare) and eluted with buffer A containing 30 mM reduced L-glutathione. The GST tag was cleaved by TEV protease overnight at 16 ℃ tag, the protein was then purified by a Superdex 75 column (GE Healthcare). Finally, proteins were concentrated to ~15 mg/ml in buffer B (50 mM $Na_2HPO_4$, 150 mM NaCl, 1 mM EDTA, 1 mM DTT, pH 7.0) and stored at -80 ℃. DMNT1 peptide was synthesized (GL Biochem), and stock solutions (5 to 15 mM) were prepared in buffer B. The peptide sequence is RRS-$K^{me1}$-SDG, in which K142 was mono-methylated.

#### Crystallization, data collection and structure determination

Crystals of apo PHF20L1 Tudor1 were grown at 16 ℃ using the sitting drop vapor diffusion method by mixing 1 μl of protein (15 mg/mL) with 1 μl reservoir buffer (10% PEG 6000, 0.1 M citric acid, PH 4.0).The PHF20L1 Tudor1 protein and DMNT1K142$^{me1}$ peptide were mixed at a 1:3 molar ratio and incubated overnight at 4 ℃. Crystals of PHF20L1 Tudor1 in complex with K142$^{me1}$ DNMT1 were grown at 16 ℃ using the hanging drop vapor diffusion method by mixing 1 μl of mix with 1 μl reservoir buffer (30% PEG MME 2000, 0.1 M sodium cacodylate, pH 6.0).

Crystals of PHF20L1 in complex with MES were grown at 16 ℃ using the sitting drop vapor diffusion method by mixing 1 μl of protein (15 mg/mL) with 1 μl reservoir buffer (20% PEG 8000, 0.2 M $(NH_4)_2SO_4$, 0.1 mM MES, PH 6.5). The PHF20L1 Tudor1 protein and hit **1** were mixed at a 1:3 molar ratio and incubated overnight at 4 ℃. Crystals of PHF20L1 Tudor1 in complex with hit **1** were grown at 16 ℃ using the hanging drop vapor diffusion method by mixing 1 μl of mix with 1 μl reservoir buffer (1.6 M magnesium sulfate, 0.1

mM MES, PH 6.5). Crystals of PHF20L1 Tudor1 Y24L and Y24W/Y29W mutants were grown at 16 ℃ using the hanging drop vapor diffusion method by mixing 1 μl of protein (15 mg/mL) with 1 μl reservoir buffer (1.6 M lithium sulfate, 0.1 M Tris, PH 8.0 and 30% PEG 4000, 0.2 M Lithium Sulfate, 0.1M Tris, PH 8.0, respectively).

Crystals were soaked in the cryo protectant made of mother liquor supplemented with 25% glycerol before being flash-frozen in liquid nitrogen. Datasets were collected on Beamline 19U1 at Shanghai Synchrotron Radiation Facility (SSRF). The structure of the free form PHF20L1 Tudor1 was solved by molecular replacement with the program MOLREP [1], using the free-form PHF20 Tudor1 (PDB ID: 3SD4) as the search model. This free form PHF20L1 Tudor1 structure was then recruited as the search model in the molecular replacement using program MOLREP for the structures of PHF20L1 Tudor1 mutants or wild-type PHF20L1 Tudor1 in complex with $K142^{me1}$ DNMT1, MES, or hit **1**. The structures were modeled in COOT and then refined by the programs REFMAC5 and PHENIX [2-4]. Crystal diffraction data and refinement statistics were displayed in Table S1 and S2. Structure analysis was performed using COOT and PyMOL (http://www.pymol.org/).

**NMR backbone chemical shift assignment**

All NMR spectra of $^{15}N/^{13}C$-labeled PHF20L1 Tudor1 (0.6 mM) in a buffer (50 mM $Na_2HPO_4$, 150 mM NaCl, 1 mM EDTA, 1 mM TCEP, pH 7.0, 10% $D_2O$) were recorded at 298 K on a Bruker DMX 600 spectrometer equipped with a cryoprobe. Sequential backbone chemical shift assignments were carried out by analysis of 3D HNCO, HNCA, CBCANH, and CBCA(CO)NH experiments. All NMR spectral data were processed with NMRPipe and NMRDraw software [5]. All assignments were generated using Sparky software (T. D. Goddard and D. G. Kneller, SPARKY 3, University of California, San Francisco).

**NMR chemical shift perturbations**

The $^{15}N$-labeled PHF20L1 Tudor1 proteins were concentrated to 50 μM in PBS buffer, which were then titrated by the ligands. The HSQC spectra were acquired on Agilent 700 MHz spectrometer at the ligand/protein molar ratio ranging from 0.0 to 4.0. The binding constant was best-fitted assuming a 1: 1 binding mode.

**Isothermal Titration Calorimetry**

ITC assays were performed on a MicroCal iTC200 calorimeter (GE Healthcare) at 25 ℃. The concentrations of proteins were determined using UV-vis spectroscopy, while the peptide concentrations were determined using quantitative NMR. Protein and peptide were dialyzed in a phosphate buffer (50 mM $Na_2HPO_4$, 150 mM NaCl, pH 7.0) and concentrated to 0.2 mM and 5 mM, respectively. Curve fitting to a single binding site model was performed by the ITC data analysis module of Origin 7.0 (MicroCal) provided by the manufacturer.

#### $^{15}$N relaxation dispersion

The $^{15}$N-labeled PHF20L1 Tudor1 proteins were prepared as described previously. All NMR relaxation dispersion spectra were acquired at 287 K on an Agilent 700 MHz or Bruker 600 MHz spectrometer equipped with cryoprobe. The $\tau_{cp}$ was defined as the delay between consecutive inversion pulses in the CPMG periods respectively. All two-dimensional data were recorded as a complex data matrix comprise of 96×88 points. The total relaxation time $T_{CPMG}$ is 40 ms. The effective $^{15}$N transverse relaxation rate $R_2^{eff}$ was calculated from the peak intensities according to the following equation,

$$R_2^{eff}(\tau_{cp}) = -\frac{1}{T_{CPMG}} Ln[\frac{I(\tau_{cp})}{I_0}] \tag{S1}$$

where $I(\tau_{cp})$ represented the peak intensity at a certain $\tau_{cp}$ delay and $I_0$ denoted the corresponding peak intensity of the reference spectrum without the CPMG pulse. The relaxation dispersion profiles were best-fitted using GLOVE program assuming a two-state exchange model.

#### Conformational selection model of ligand binding

The free-form PHF20L1 Tudor1 was in equilibrium between the closed (denoted $P$) and open state ($P^*$) as follows,

$$P \leftrightarrow P^* \tag{S2}$$

where ρ denoted the population of the open state. The ligand ($L$) binding was described as below,

$$P^* + L \leftrightarrow PL \tag{S3}$$

The dissociation constant $K_D^*$ of this open state was defined as $[P^*][L]/[PL]$, which represented the concentration of $P^*$, $L$ and the complex form $PL$, respectively. The apparent $K_D^{app}$, defined as $([P]+[P^*])[L]/[PL]$, was thus derived as,

$$K_D^{app} = K_D^*/\rho \tag{S4}$$

That is to say, the experimentally determined affinity can be mediated by the population of the open state reciprocally.

**Molecular Dynamics (MD) simulations**

MD simulations of the wild-type, Y24L, and Y24W/Y29W mutants were performed using the AMBER16 software package (San Francisco, CA, USA). The simulated system was built in the tleap module using the ff14SB force field [6]. The structure was immersed into a truncated octahedral box that extended 10 Å away from the solute border, using the TIP3P water model and periodic boundary conditions [7]. The total numbers of atoms for the three systems were 17480 for close-form, 12892 for Y20L, and 16230 for Y24W/Y29W with reaching a salt concentration of 150mM NaCl.To remove the bad contacts,the waters and ions were initially minimized for 2000 steps using the steepest descent method for the first 1000 steps and then the conjugate gradient algorithm for the last 1000 steps, with the position of protein fixed (force constant was 500 kcal $mol^{-1}$ $Å^{-2}$). In the second energy minimization stage, the restraints on the protein were removed. This stage was conducted for 2500 steps, using the steepest descent method in the first 1000 steps and then the conjugate gradient algorithm for the last 1500 steps. After that, a heat-up MD was run at a constant volume. The system was heated from 0 to 287 K for 100 ps with a weak restraint of 10 kcal $mol^{-1}$ $Å^{-2}$ on the solute. Then, free MD simulations of 1μs were carried out under the NPT condition utilizing the GPU accelerated pmemd.cuda code.Temperature was regulated using the Langevin dynamics with a collision frequency of 1.0 $ps^{-1}$ [8]. Pressure was controlled with isotropic position scaling at 1 bar with a relaxation time of 2.0 ps. All of the bonds involving hydrogen atoms were constrained using the SHAKE algorithm [9]. A 2 fs integration step was used. The long-range electrostatic interactions were calculated using PME method with a 10 Å cutoff for the range-limited non-bonded interactions [10].

**Principal Component Analysis (PCA)**

In a PCA, the correlated internal motion with N degrees of freedom can be described by the covariance matrix,

$$\sigma_{mn} = \langle (r_m - \langle r_m \rangle)(r_n - \langle r_n \rangle) \rangle \quad \text{(S5)}$$

Where $r_1$ , ⋯; $r_N$ mean the input coordinates and $\langle \cdots \rangle$ represents the average over all sampled conformations. Diagonalization of this covariance matrix results in N eigenvectors $\{V^{(i)}\}$ and eigenvalues $\{\lambda_i\}$, which describe the modes of the collective motion and their respective amplitudes. The PCs are the projections of the coordinates $r$ onto the eigenvectors.

$$\chi_i = V^{(i)} \cdot r \quad \text{(S6)}$$

There are numerous options to define input coordinates to perform a PCA. Here we use both the heavy atoms of residues and contact distance between residues as input coordinates.

**Contact-Based Principal Component Analysis (ConPCA)**

We consider a contact as formed with distance between closest lying non-hydrogen atoms of two residues $i$ and $j$ [11]. First the contacts of closed, mutants of Y24L, Y24W/Y29W need to be determined. We did 1 μs MD simulation of these three systems from the crystal structures and considered seven residue-residue contacts of D23, Y24, L25, Y29, F47, W50 and Y54. We got total 21 pairs residue-residue contacts and calculated the contact distance $D_{ij}$ of all identified contacts for all frames of the 1 μs MD trajectory by g_mindist tools of Gromacs program [12]. Then this data of distance is used as input for the ConPCA. Employing this definition, we calculate the covariance matrix,

$$\sigma_{\mu\upsilon} = \langle (D_\mu - \langle D_\mu \rangle)(D_\upsilon - \langle D_\upsilon \rangle) \rangle \quad \text{(S7)}$$

which defines the conPCA.

**Pull-down assay**

For pull-down assay of the biotin-labeled $K142^{me1}$DNMT1-peptide against the GST-PHF20L1-Tudor1, the high capacity streptavidin agarose resins (Thermo) with about 50 μg biotin-labeled $K142^{me1}$DNMT1-peptide were incubated with GST-PHF20L1 Tudor1 (100 μg) upon the titration of hit **1** in a buffer containing 50 mM $Na_2HPO_4$, 150 mM NaCl, 1 mM EDTA, 1 mM DTT, pH 7.0 for 2 h at 4 ℃. The molar ratio of hit **1** :

GST-Tudor1 were 100, 50, 25, and 12.5, respectively. The resins were pelleted and washed five times using the same buffer. The captured proteins were eluted and analyzed using SDS-PAGE.

**Immunostaining and fluorescence colocalization**

Cells were cultured in 24-well chamber slides and transfected with recombinant plasmids of either wild type (WT) or RFP (Y1421A/N1464A) chimeras of PHF20L1. After washing twice with PBS, cells were fixed in 4% paraformaldehyde and permeabilized in PBS containing 0.5% Triton X-100 for 5 min at room temperature. After blocking with 5% BSA in PBS for 1 h, cells were incubated with primary antibodies and goat anti-rabbit immunoglobulin G conjugated with FITC antibody (invitrogen), respectively. DAPI (Sigma) was used for nuclear staining. After extensive washing, cells were analyzed by a Zeiss LSM880 laser confocal system (Zeiss).

**Coimmunoprecipitation**

The 293T cells were simultaneously transfected with 3 μg of GFP-DNMT1 and 2 μg of Flag-PHF20L1 (WT or mutants) plasmids using Lipofectamine 3000 (Invitrogen). Seven 6-cm dishes of 293T cells were harvested, washed with PBS. Cell extracts were prepared in modified RIPA buffer (50 mM Tris, pH 7.4, 150 mM NaCl, 1 mM EDTA, 1% NP40, 1 mM protease inhibitor cocktails) and incubated with anti-GFP affinity agarose (Chromotek) with gentle shaking overnight at 4 ℃. The resin was washed using modified RIPA buffer. Eluted proteins were then separated by SDS-PAGE and electroblotted to nitrocellulose membranes. Membranes were blocked in 1× PBS-T (0.1% tween-20) and fat-free dry milk or 5% bovine serum albumin in blocking buffer for 1 h at room temperature. Membranes were incubated with the primary antibodies diluted in the blocking buffer overnight at 4 ℃. Membranes were washed three times in 1× PBS-T (0.1% tween-20), 10 min each. Secondary antibodies conjugated with horseradish peroxidase were added, and the membranes were incubated for 1 h at room temperature. Membranes were washed three times in1× PBS-T (0.1% tween-20) for 15-20 min each. ECL Western blotting reagent kit (ThermoScientific) was used for protein detection.

**SI References**


1. Vagin, A. and A. Teplyakov, *Molecular replacement with MOLREP.* Acta Crystallographica Section D-Biological Crystallography, 2010. **66**: p. 22-25.
2. Adams, P.D., et al., *PHENIX: a comprehensive Python-based system for macromolecular structure solution.* Acta Crystallographica Section D-Structural Biology, 2010. **66**: p. 213-221.

3. Emsley, P., et al., *Features and development of Coot.* Acta Crystallographica Section D-Biological Crystallography, 2010. **66**: p. 486-501.

4. Murshudov, G.N., A.A. Vagin, and E.J. Dodson, *Refinement of macromolecular structures by the maximum-likelihood method.* Acta Crystallographica Section D-Structural Biology, 1997. **53**: p. 240-255.

5. Delaglio, F., et al., *Nmrpipe - a Multidimensional Spectral Processing System Based on Unix Pipes.* Journal Of Biomolecular Nmr, 1995. **6**(3): p. 277-293.

6. Maier, J.A., et al., *ff14SB: Improving the Accuracy of Protein Side Chain and Backbone Parameters from ff99SB.* Journal Of Chemical Theory And Computation, 2015. **11**(8): p. 3696-3713.

7. Mark, P. and L. Nilsson, *Structure and dynamics of the TIP3P, SPC, and SPC/E water models at 298 K.* Journal Of Physical Chemistry B, 2001. **105**(43): p. 24a-24a.

8. Pastor, R.W., B.R. Brooks, and A. Szabo, *An Analysis Of the Accuracy Of Langevin And Molecular-Dynamics Algorithms.* Molecular Physics, 1988. **65**(6): p. 1409-1419.

9. Forester, T.R. and W. Smith, *SHAKE, rattle, and roll: Efficient constraint algorithms for linked rigid bodies.* Journal Of Computational Chemistry, 1998. **19**(1): p. 102-111.

10. Darden, T., D. York, and L. Pedersen, *Particle Mesh Ewald - an N.Log(N) Method for Ewald Sums In Large Systems.* Journal Of Chemical Physics, 1993. **98**(12): p. 10089-10092.

11. Ernst, M., F. Sittel, and G. Stock, *Contact- and distance-based principal component analysis of protein dynamics.* Journal Of Chemical Physics, 2015. **143**(24).

12. Pronk, S., et al., *GROMACS 4.5: a high-throughput and highly parallel open source molecular simulation toolkit.* Bioinformatics, 2013. **29**(7): p. 845-854.

**Table S1.** Crystallography data collection and refinement statistics of PHF20L1 Tudor1 and its complexes.

| **Data collection** | Apo-PHF20L1Tudor1 | +DNMT1 | +MES | +hit **1** |
|---|---|---|---|---|
| Beamline | 19U, SSRF | 19U, SSRF | 19U, SSRF | 19U, SSRF |
| Space group | P1 | $P2_12_12$ | $P2_12_12_1$ | $P2_12_12_1$ |
| Wavelength (Å) | 0.9789 | 0.9789 | 0.9789 | 0.9789 |
| Resolution (Å) | 25.32-1.30 (1.35-1.30)[a] | 33.24-1.90 (1.93-1.90)[a] | 37.10-1.60 (1.63-1.60)[a] | 36.90-1.23 (1.28-1.23)[a] |
| Cell dimensions | | | | |
| a, b, c (Å) | 25.35 32.02 47.35 | 42.70 52.93 31.39 | 52.18 52.77 101.38 | 51.80 52.57 101.39 |
| α, β, γ (°) | 84.99 88.22 82.85 | 90 90 90 | 90 90 90 | 90 90 90 |
| Unique reflections | 29808 (1817) | 5921 (561) | 37615 (1897) | 80879 (7954) |
| Completeness (%) | 89.3 (82.3) | 99.0 (93.7) | 99.8 (99.6) | 100 (99.9) |
| Redundancy | 3.0 (2.5) | 6.3 (4.8) | 9.4 (8.5) | 8.9 (9.0) |
| I/σI | 21.2 (2.3) | 16.9 (2.1) | 24.8 (3.5) | 33.5 (7.1) |
| $R_{merge}$ (%) | 4.9 (36.3) | 11.9 (54.1) | 7.3 (42.0) | 6.4 (25.1) |
| **Refinement** | | | | |
| $R_{work}$ (%) | 15.25 | 19.22 | 19.41 | 18.36 |
| $R_{free}$ (%) | 18.18 | 22.97 | 22.09 | 20.03 |
| No. of atoms | | | | |
| Protein | 1201 | 590 | 2318 | 2401 |
| Ligand | | 45 | 48 | 56 |
| Water | 200 | 32 | 331 | 441 |
| Average B factors (Å$^2$) | | | | |
| Protein | 17.35 | 25.66 | 21.76 | 12.32 |
| Ligand | | 45.25 | 43.81 | 13.14 |
| Water | 31.27 | 32.35 | 25.79 | 20.31 |
| Root mean square deviations | | | | |
| Bond lengths (Å) | 0.005 | 0.006 | 0.006 | 0.005 |
| Bond angles (°) | 0.93 | 0.84 | 0.92 | 1.18 |
| Ramachandran plot | | | | |
| Favored (%) | 98.5 | 100 | 97.4 | 97.0 |
| Allowed (%) | 1.5 | 0 | 2.6 | 3.0 |
| Disallowed | 0 | 0 | 0 | 0 |

[a] Values for the highest-resolution shell are shown in parentheses.

**Table S2.** Binding affinities of hit **1** and K142$^{me1}$ DNMT1 peptide to various mutants of PHF20L1 Tudor1.

| PHF20L1 Tudor1 | +hit **1** | + K142$^{me1}$ DNMT1 peptide |
|---|---|---|
| | $K_D$ (mM)[a] | $K_D$ (mM)[a] |
| Wild-type | 0.31 ± 0.05[b] | 0.67 ± 0.08 |
| Y24A | 0.30 ± 0.05 | |
| Y24L | 0.30 ± 0.04 | 0.4 ± 0.2 |
| Y24W | 1.5 ± 0.4 | ND[c] |
| Y29W | 0.41 ± 0.06 | ND |
| Y24L/Y29L | 0.53 ± 0.07 | |
| Y24W/Y29W | 2.3 ± 0.08 | ND |
| R49G | 0.18 ± 0.01 | |
| R49W | 0.22 ± 0.02 | |
| W50A | ND | ND |
| Y54A | ND | ND |

[a] Dissociation constants ($K_D$) determined by the dose-dependent chemical shift perturbation of $^{15}$N-labeled PHF20L1Tudor1 upon titration of hit **1**. [b] Fitting error throughout the table. [c] No detectable chemical shift perturbations.

**Table S3.** Crystallography data collection and refinement statistics of PHF20L1 Tudor1 mutants.

| **Data collection** | Y24L_mutant | Y24W/Y29W_mutant |
|---|---|---|
| Beamline | 19U, SSRF | 19U, SSRF |
| Space group | $P4_1$ | $P6_5$ |
| Wavelength (Å) | 0.9785 | 0.9785 |
| Resolution (Å) | 37.17-1.58 (1.61-1.58)[a] | 27.33-1.85 (1.88-1.85)[a] |
| Cell dimensions | | |
| a, b, c (Å) | 52.57 52.57 38.65 | 63.12 63.12 26.98 |
| α, β, γ (°) | 90 90 90 | 90 90 120 |
| Unique reflections | 13228 (1319) | 5374 (522) |
| Completeness (%) | 100 (99.9) | 100 (99.5) |
| Redundancy | 13.2 (12.1) | 9.9 (8.3) |
| I/σI | 37.4 (6.3) | 25.0 (2.7) |
| $R_{merge}$ (%) | 9.7 (34.1) | 10.1 (69.6) |
| **Refinement** | | |
| $R_{work}$ (%) | 17.92 | 16.01 |
| $R_{free}$ (%) | 19.09 | 20.77 |
| No. of atoms | | |
| Protein | 605 | 585 |
| Ligand | | |
| Water | 35 | 81 |
| Average B factors (Å$^2$) | | |
| Protein | 18.44 | 25.95 |
| Ligand | | |
| Water | 32.78 | 32.04 |
| Root mean square deviations | | |
| Bond lengths (Å) | 0.003 | 0.004 |
| Bond angles (°) | 1.16 | 0.67 |
| Ramachandran plot | | |
| Favored (%) | 100 | 98.5 |
| Allowed (%) | 0 | 1.5 |
| Disallowed | 0 | 0 |

[a] Values in parentheses are for the highest-resolution shell.

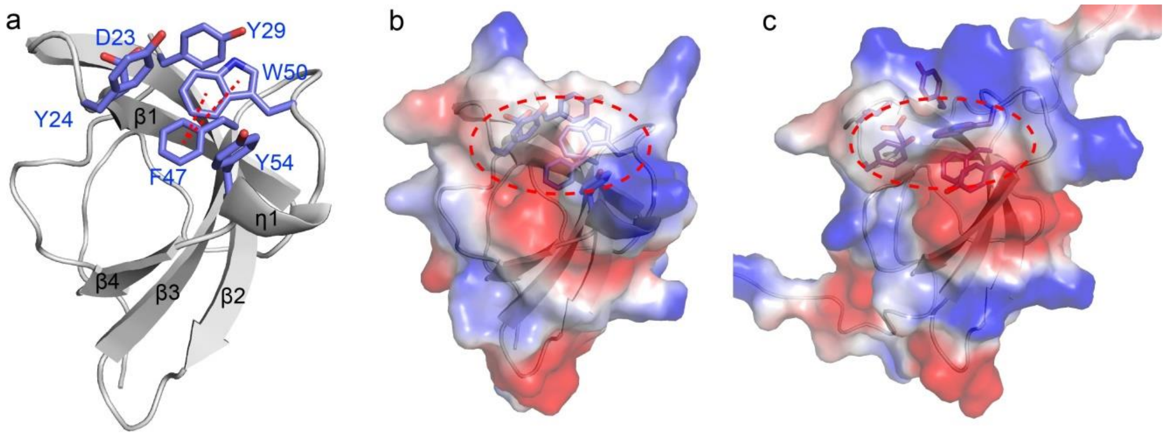


**Figure S1.** The crystal structure of the free form PHF20L1 Tudor1. **a**) The cartoon view of PHF20L1 Tudor1. The aromatic cage residues are annotated and highlighted as sticks, and the dotted lines denote π-π interactions. **b**) The electrostatic surface representation of PHF20L1 Tudor1, where the blocked aromatic cage is highlighted in the circle. **c**) The electrostatic surface representation of the free-form solution structure of PHF20L1 Tudor1 (PDB code: 2EQM)

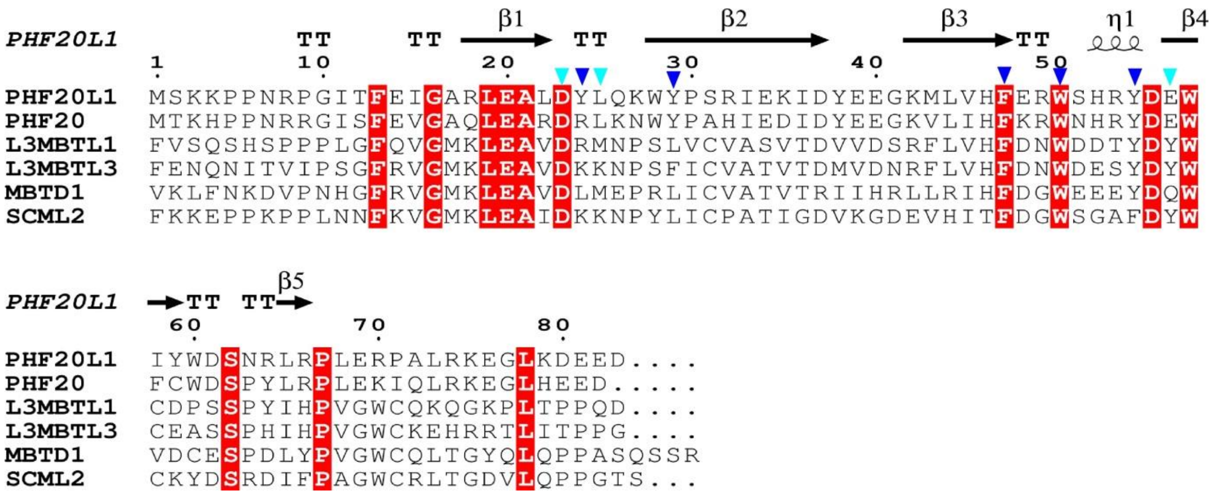


**Figure S2.** Sequence blast of PHF20L1 Tudor1 in the protein data bank. These of highest sequence identities are Tudor and MBT domains, which belong to the Royal superfamily. Marked are residues proximal to the aromatic cage, in which the aromatic residues are annotated by blue triangles.

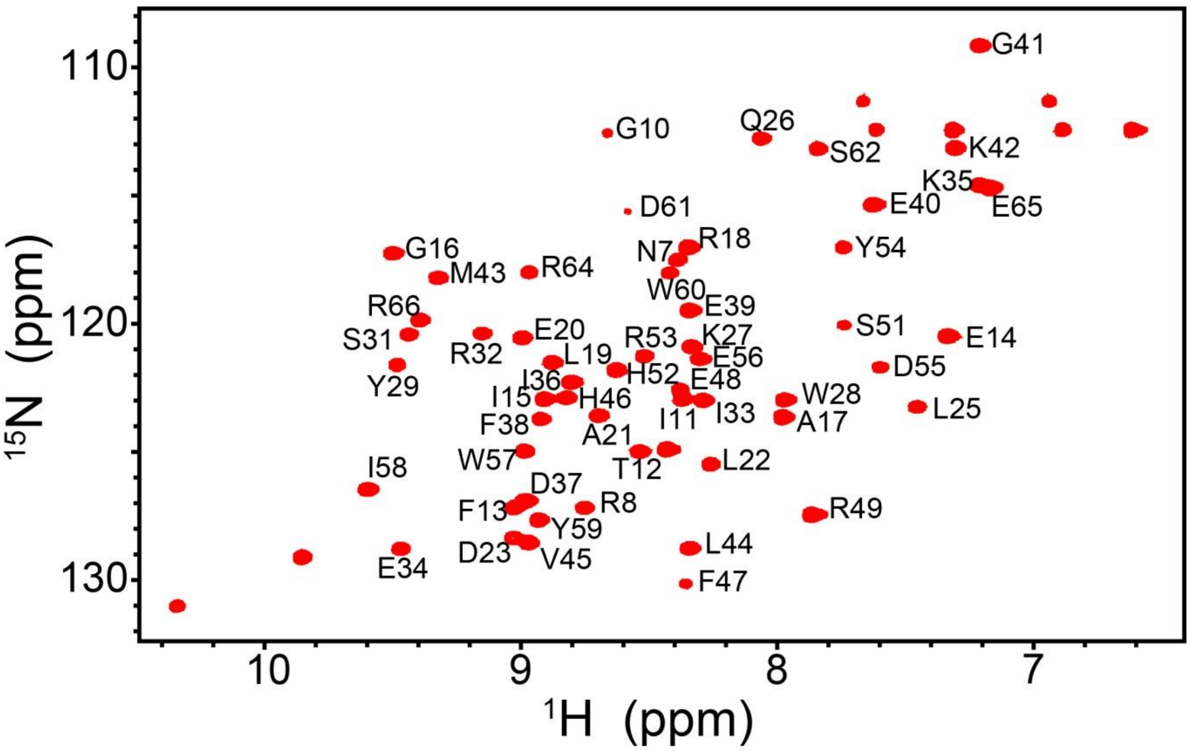


**Figure S3.** Backbone $^{1}H$-$^{15}N$ chemical shift assignment of PHF20L1 Tudor1.

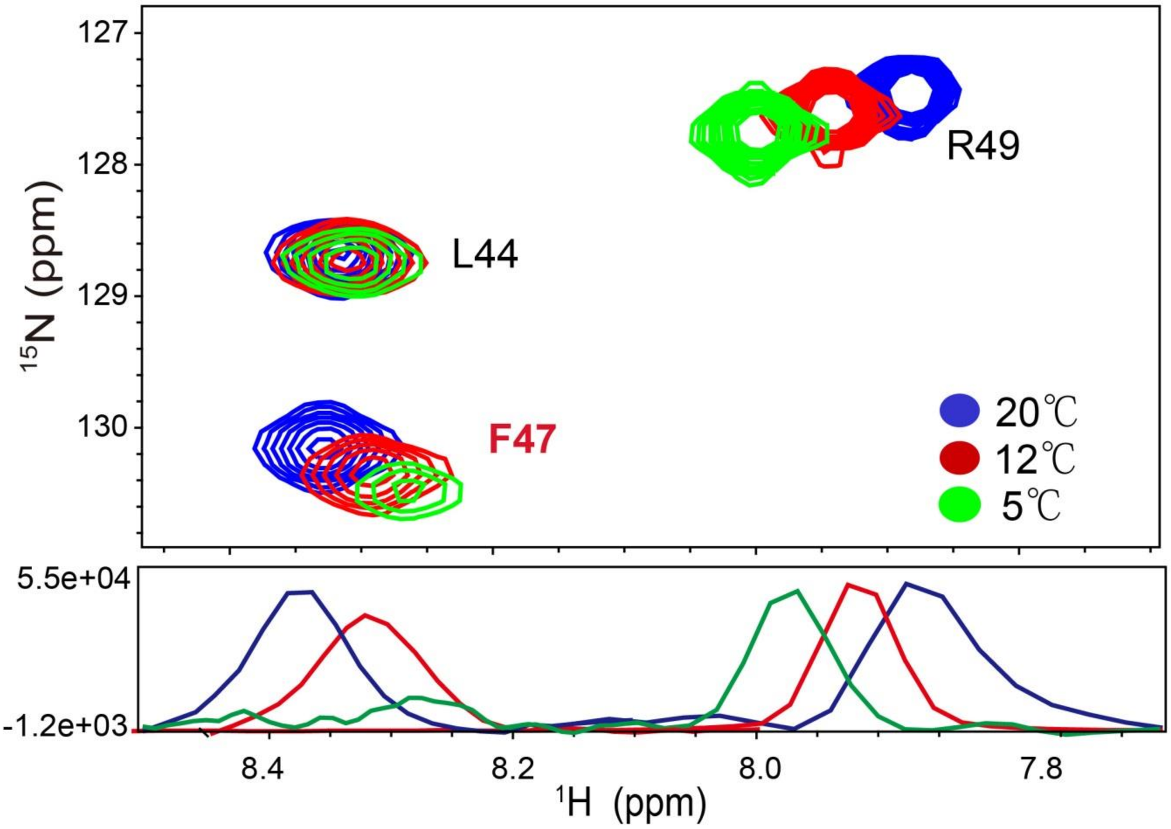


**Figure S4.** The representative signals in the $^{1}$H-$^{15}$N HSQC spectra of PHF20L1 Tudor1 acquired at variable temperatures.

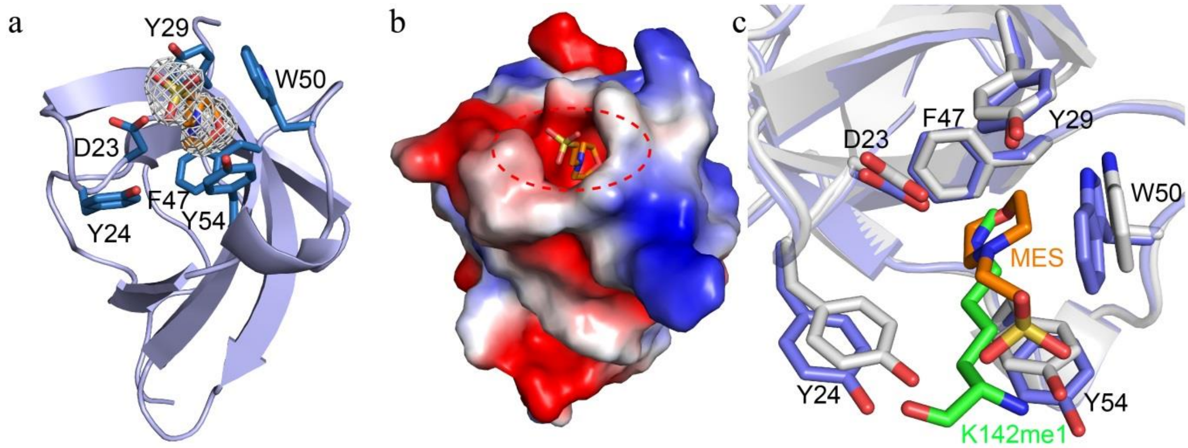


**Figure S5.** MES binds to the open form aromatic cage of PHF20L1 Tudor1. **a**) Cartoon representation of PHF20L1 Tudor1 (slate) in complex with MES (orange), whose 2Fo-Fc electron-density map is contoured at 1.0 σ (gray). Residues in the aromatic cage (sticks) adopt the open form to accommodate MES. **b**) The electrostatic surface of PHF20L1 Tudor1 in complex with MES (sticks). The open form aromatic cage of PHF20L1 Tudor1 is highlighted in the red oval. **c**) Superimposition of the crystal structures of PHF20L1 Tudor1 in complex with MES (sticks in orange) and K142$^{me1}$ DNMT1 (sticks in green). The aromatic cage residues (sticks in slate and gray) adopt similar conformations in the two complex structures.

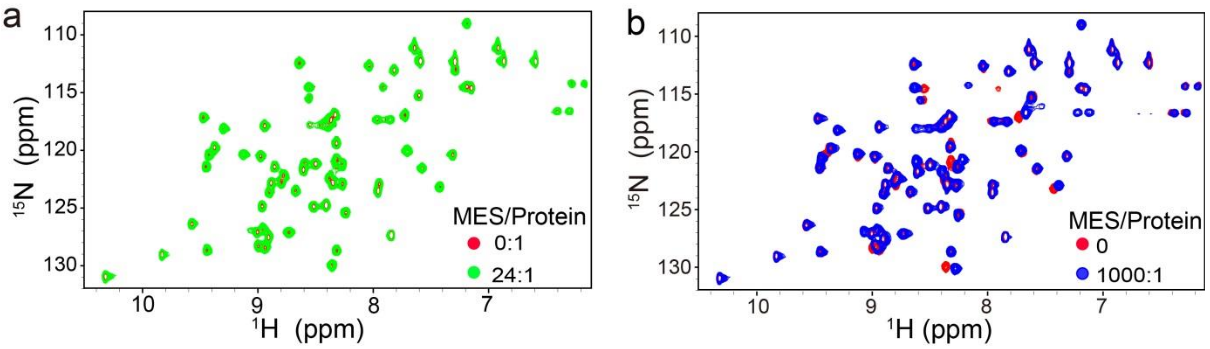


**Figure S6**. The$^{1}$H-$^{15}$N HSQC spectra of PHF20L1 Tudor1 superimposed at different MES/protein molar ratios.

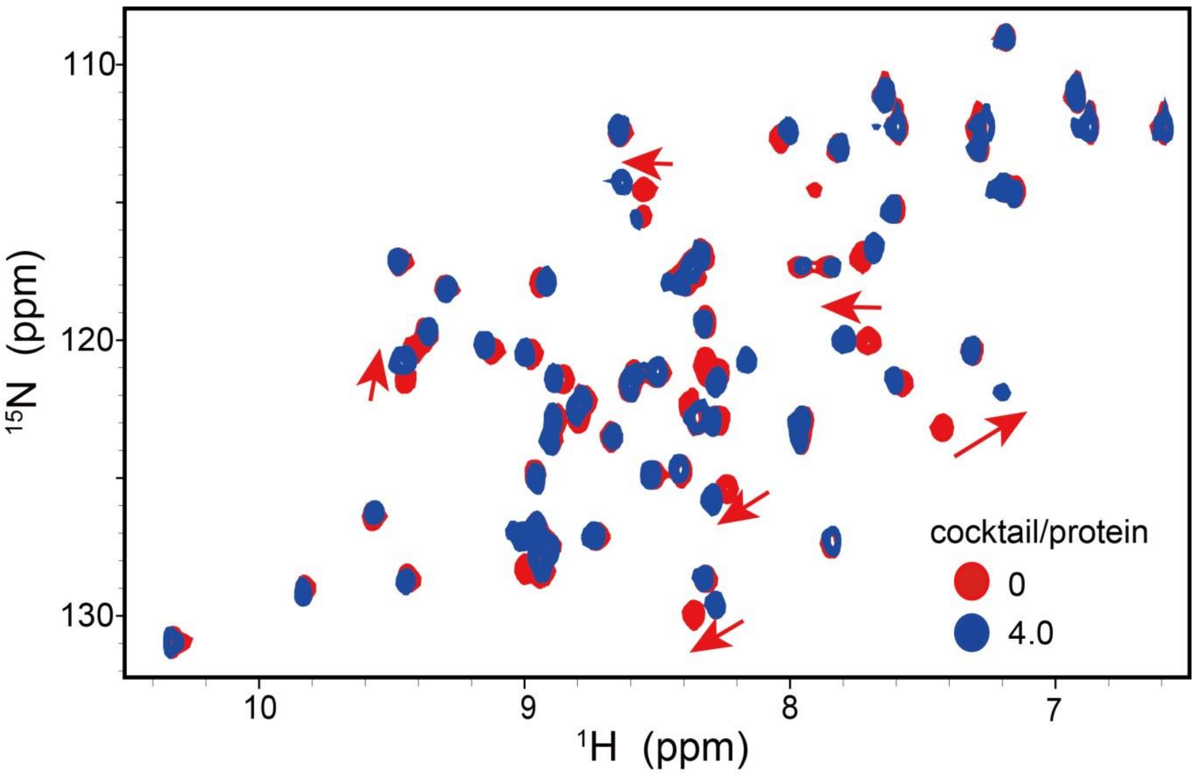


**Figure S7**. Protein-observed fragment-based screening against $^{15}$N-labeled PHF20L1 Tudor1. Superimposed are the $^{1}$H-$^{15}$N HSQC spectra at two cocktail/protein molar ratios.

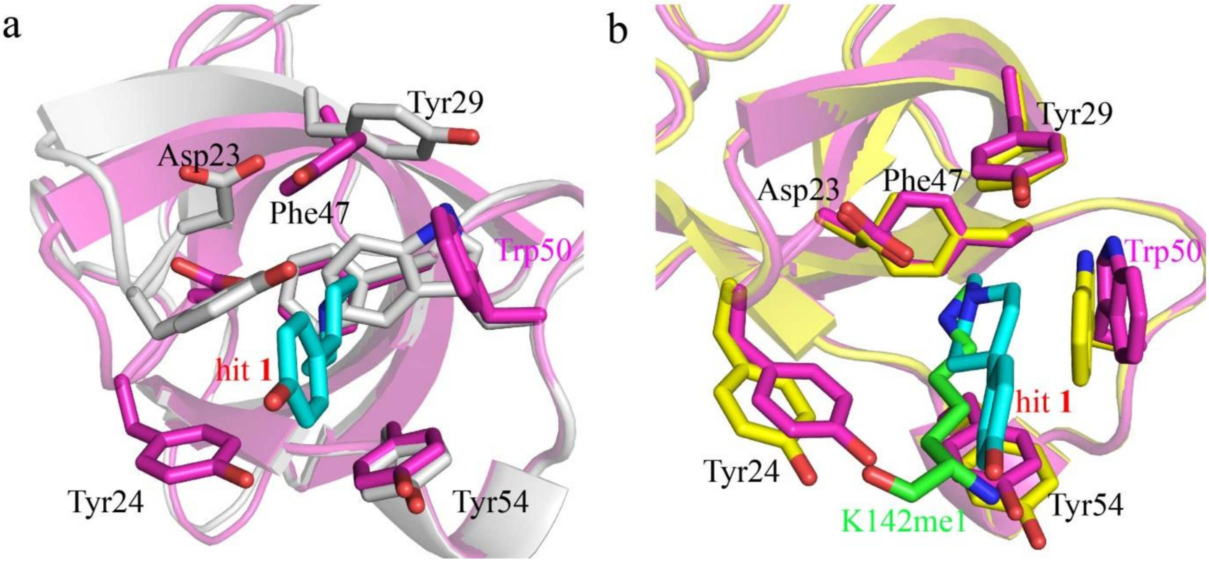


**Figure S8.** Structural comparison of the crystal structures of PHF20L1 Tudor1 and its complexes. **a**) Superimposition of the apo-PHF20L1 Tudor1 (grey) and its complex with hit **1** (Magenta). Hit **1** is shown in sticks with carbon atoms in cyan. **b**) Superimposition of the crystal structures of PHF20L1 Tudor1 in complex with hit **1** (Protein in magenta, hit **1** in cyan) and DNMT1 K142me1 peptide (protein in yellow, $K142^{me1}$ in green).

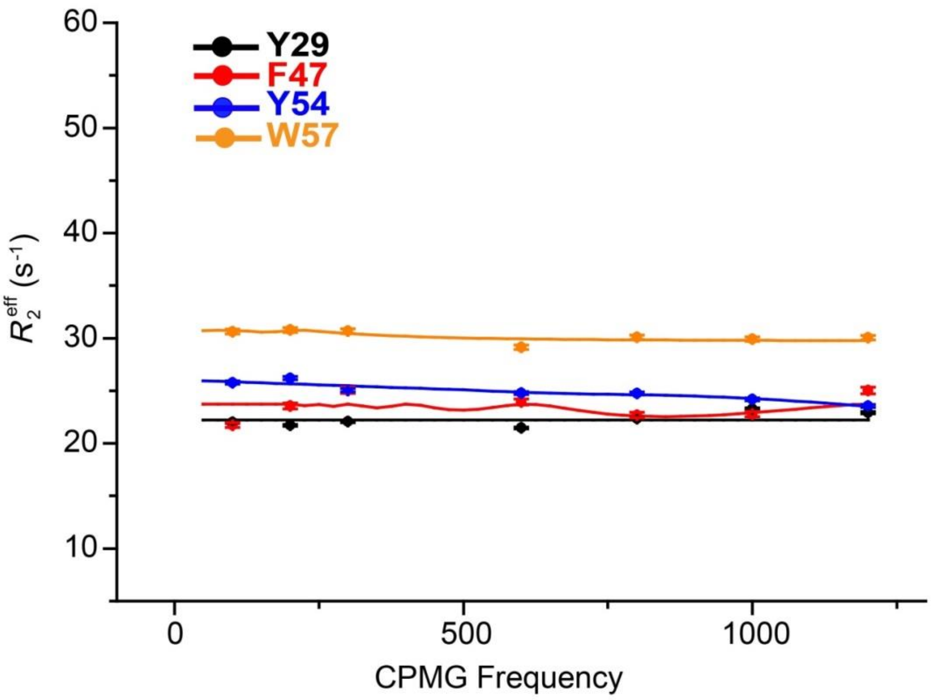


**Figure S9.** Relaxation dispersion profiles for typical residues of PHF20L1 Tudor1 in the presence of 8-fold excess of K142$^{me1}$ DNMT1 peptide.

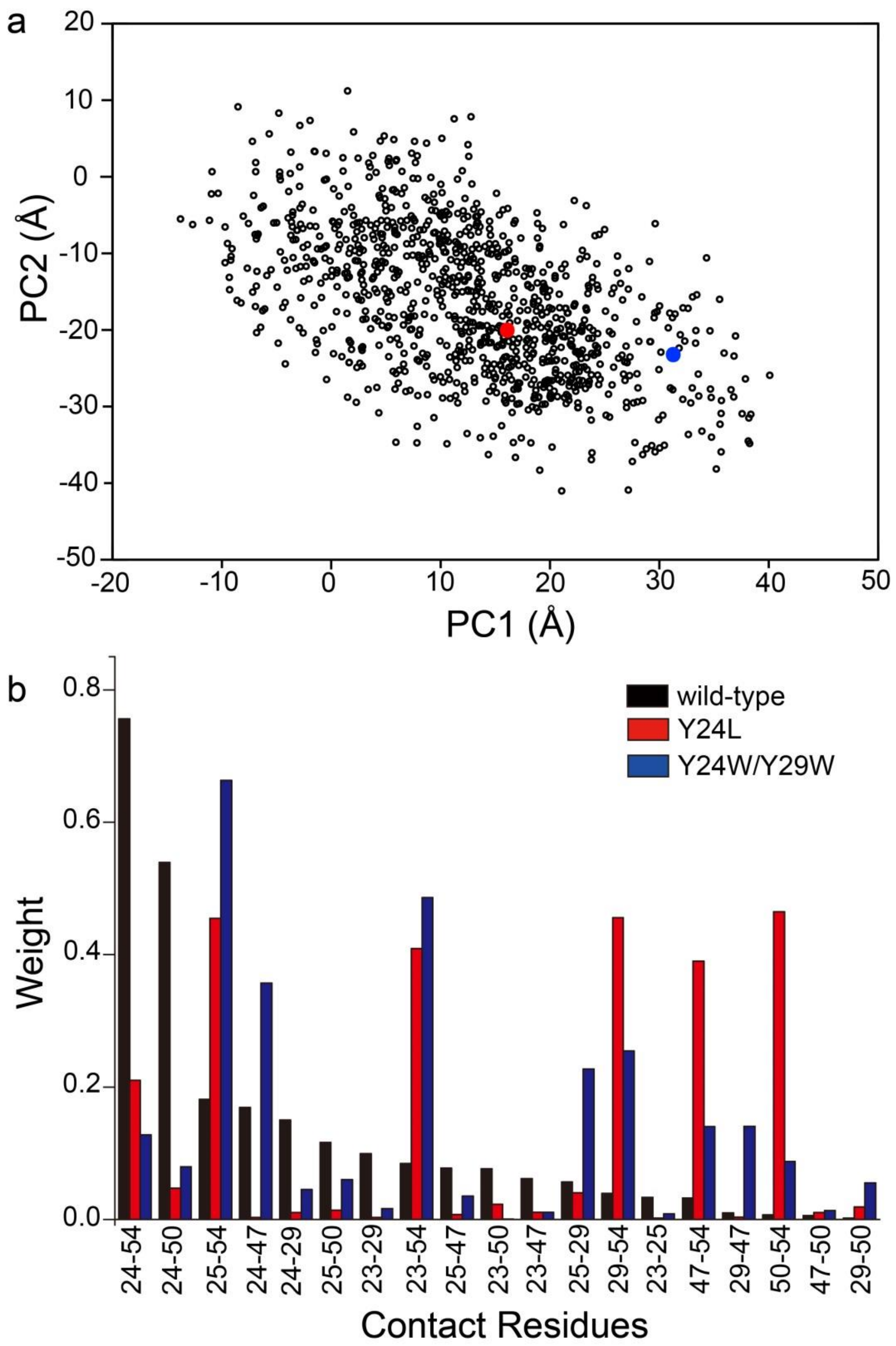


**Figure S10. a**) Projection of the trajectory of closed state onto the 2D essential subspace defined by the PC1 and PC2. PCA was performed on the 1004 simulated structures of the WT and each point on the plane represents a conformation. The 1004 simulated structures are colored black. The projection of crystal structures of WT and open states colored by red and blue, respectively. **b**) Contact-based PCA. Elements of the normalized first eigenvector $\{\upsilon_i\}$ of conPCA, with index $i$ labeling the considered contacts of closed states, Y24L and Y24W/Y29W mutants. Only components that constitute up to 90% of the norm are shown.

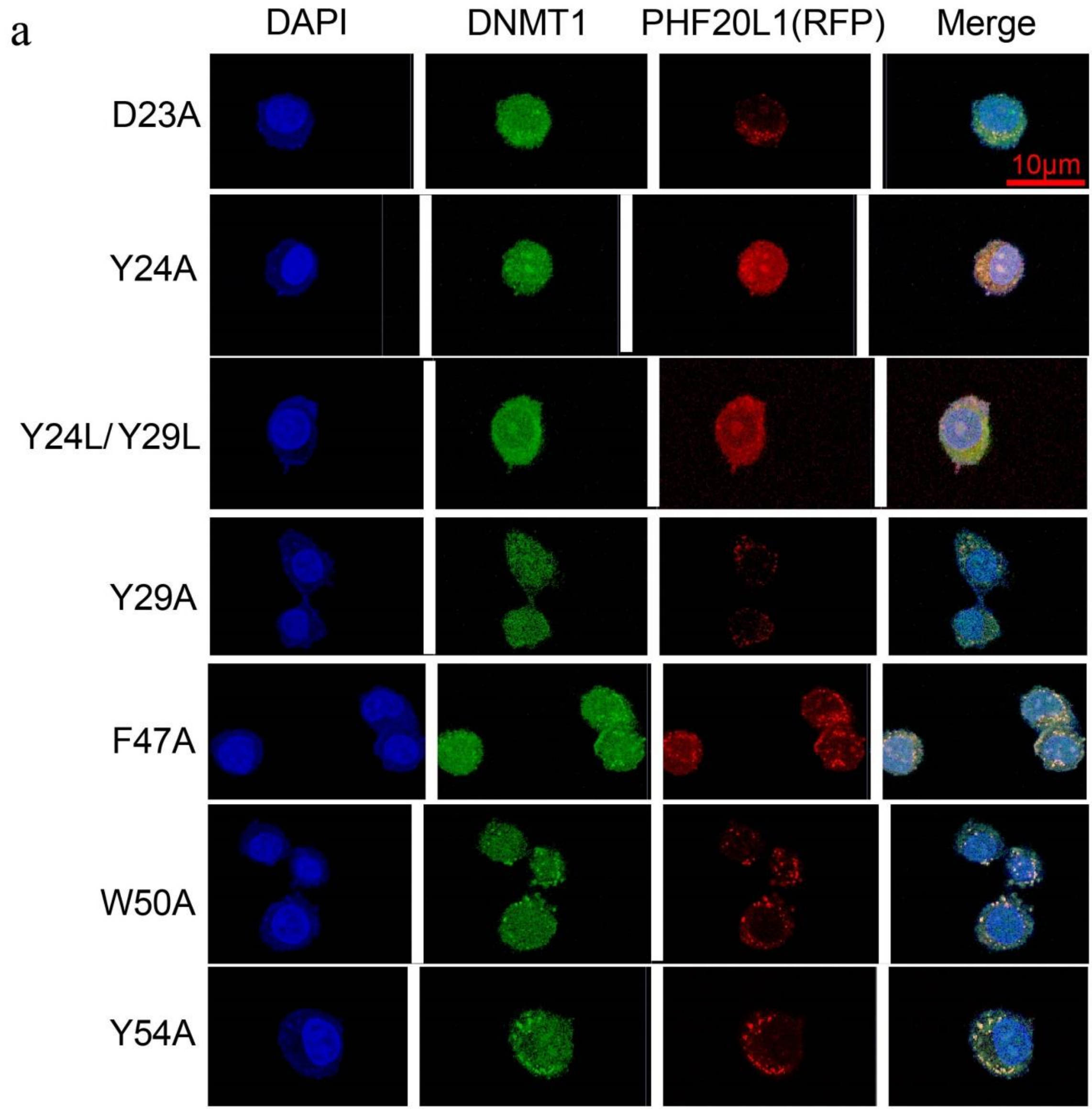


**Figure S11**. The immunofluorescence assay of endogenous DNMT1(green stained by antibodies) and ectopically over-expressed RFP-fused PHF20L1 in Hela cells.
